# A Ventromedial Hypothalamic Neuron Subset Encodes a Conspecific-Tuned Behavior State Driving Social Investigation

**DOI:** 10.1101/2025.05.30.650902

**Authors:** Shih-Che Lin, Ju-Yu Hsu, Ting-An Su, Han-Zhuan Lee, Shi-Bing Yang

## Abstract

The VMH^SF1^ neurons encode a predator-orientated internal state that drives innate defensive responses. Although neuroanatomical studies show that male mouse VMH^SF1^ neurons are interconnected with structures in the social behavioral network, how VMH^SF1^ neurons represent stimulus and behaviors under social context remain elusive. To address the question, we employed fiber photometry and miniscope Ca^2+^ imaging on VMH^SF1^ neurons and found that VMH^SF1^ neurons are strongly responsive to social stimuli with a male-biased sex preference, which requires pheromonal signals and intact BNST-VMH pathway. During social interaction, VMH^SF1^ neurons are highly tuned to investigative yet strongly silenced by consummatory social behaviors. Notably, VMH^SF1^ neurons that encode defensive actions and those encoding social investigation are two distinct subpopulations. Lastly, silencing VMH^SF1^ neurons suppresses social investigation. Altogether, our results suggest the existence of a novel, non-defense driving VMH^SF1^ neurons subset that encodes conspecific social cues with sex-bias and prompts social investigation of male mice.

**Highlights:**

1. A subset of VMH^SF1^ neurons exhibit preferential response to social cues
2. Conspecific sex bias of VMH^SF1^ neurons requires pheromonal signal and BNST input
3. Social-tuned VMH^SF1^ neurons functionally encode social investigation
4. Predator defense and social investigation recruits distinct VMH^SF1^ subgroups

## Introduction

Animal expresses a wide variety of innate social behaviors upon conspecific encounters. Proper choice of behavioral action based on acquired sensory input is critical for animal’s survival and reproduction, as this aids seizing mating opportunity, securing resources, and minimizing injury. Regulation of social behaviors involves a widespread neural network, with hypothalamus being the primary hub for behavioral decision^30,73^.

The ventromedial nucleus of hypothalamus (VMH) integrates information from sensory processing structures and relays to the downstream midbrain structure, triggering various stereotypical social behaviors^12,73^, making it a pivotal node in social circuit. The VMH neurons are composed of multiple subpopulations with unique molecular and functional features^17,39^: The estrogen receptor *α* -(ESR*α*) expressing neurons in the ventrolateral division (VMHvl) are responsible for driving most of the VMH-mediated social behaviors^32,36,44,46,48,63,70,72^. Another major neural subgroup occupying the dorsomedial/central VMH (VMHdm/c) expresses steroidogenic factor-1 (SF1, alternatively known as NR5a1)^14^. The SF1-expessing neurons in VMHdm/c, henceforth referred to as VMH^SF1^ neurons, are essential for predator-triggered defensive responses. Loss-of-function treatments on VMH^SF1^ neurons augment innate defensive behaviors, rendering animal less anxious toward predator^37,43,69,71^, whereas stimulating VMH^SF1^ neurons faithfully induces fear-associated behaviors and anxiety-like state^21,37,43,71^. Furthermore, imaging studies found that VMH^SF1^ neurons are activated by predatory cues and their activities represent diverse features of defensive behavioral state^16,37,66^.

While VMH^SF1^ neurons have been extensively studied for innate predatory defense, their roles in social behavior are much less understood. Neuroanatomical evidence suggests that VMH^SF1^ neurons are structurally interconnected with multiple regions in the social behavioral network^12,73^, including the medial preoptic area (MPOA), bed nucleus of stria terminalis (BNST), medial amygdala (MeA), and periaqueductal gray (PAG).^43,47,58,67,77^ Moreover, connections exist between VMH^SF1^ neurons and putative ESR*α* neurons in VMHvl^47,58^. These structural features likely enable VMH^SF1^ neurons to modulate social behaviors. In support of this notion, VMH^SF1^ neurons of male mouse are activated by conspecific presentation^37^, and silencing VMH^SF1^ neurons diminishes pheromone-induced enhancement of sexual behaviors in female mouse^33^. Nevertheless, the role of VMH^SF1^ neurons of male mouse in social behaviors remains elusive.

Here, we adopted fiber photometry and head-mount microendoscope imaging to monitor VMH^SF1^ neuronal activities in freely behaving male mice. Our results showed that the VMH^SF1^ neurons are responsive to social stimuli with male-favoring conspecific sex preference. We further identified conspecific-emitted pheromone, and BNST input, as vital sensory and circuit components for maintaining male-biased population response of VMH^SF1^ neurons, respectively. Moreover, social cues recruit a specific VMH^SF1^ neural ensemble, and such conspecific-responsive VMH^SF1^ neurons are separable from those tuned to predatory cue, implying that social and predatory stimuli recruit distinct VMH^SF1^subgroups. Furthermore, the social-responsive VMH^SF1^ neurons are strongly tuned to investigation during social interaction, and these neurons are silenced by consummatory or defensive social behaviors, indicating their preference to appetitive social behaviors. Finally, chemogenetic silencing VMH^SF1^ neurons render the mouse less engaged in social investigation. These results provide a neural circuit mechanism of VMH^SF1^ neurons in encoding social cues and controlling social investigation, a vital step for subsequent behavioral decisions.

## Results

### VMH^SF1^ neural population is robustly activated by social cue with male-biased conspecific sex preference

We started by addressing how VMH^SF1^ neurons respond to social and non-social cues using fiber photometry in VMH^SF1^ neurons (Figure 1A-B). We implemented a simple stimulus exposure test, in which multiple categories of external stimuli were sequentially introduced with randomized order into the test animal’s home cage (Figure 1C). Consistent with previous study^37^, we found that compared to neutral non-social stimulus, encountering a novel mouse triggered a more rapid-onset, robust, and persistent population response in VMH^SF1^ neurons (Figure 1D-E). However, we also noticed that encountering a male mouse evoked substantially stronger responses than a female mouse (Figure 1D-F). The male-biased population response could either result from the same group of neurons showing distinct responsiveness to different conspecific sexes or the recruitment of additional male-responsive neurons. To distinguish the alternatives, we investigated stimulus-induced neural dynamics at cellular resolution by performing microendoscopic Ca^2+^ imaging (Figure 1G-I). We found that conspecific cues recruited more VMH^SF1^ neurons than neutral objects, and more male-responsive neurons were identified than female-responsive neurons (Figure 1J-M); indicating that an increase in stimulus-triggered VMH^SF1^ population response likely reflects the recruitment of additional neural subset.

**Figure 1.**
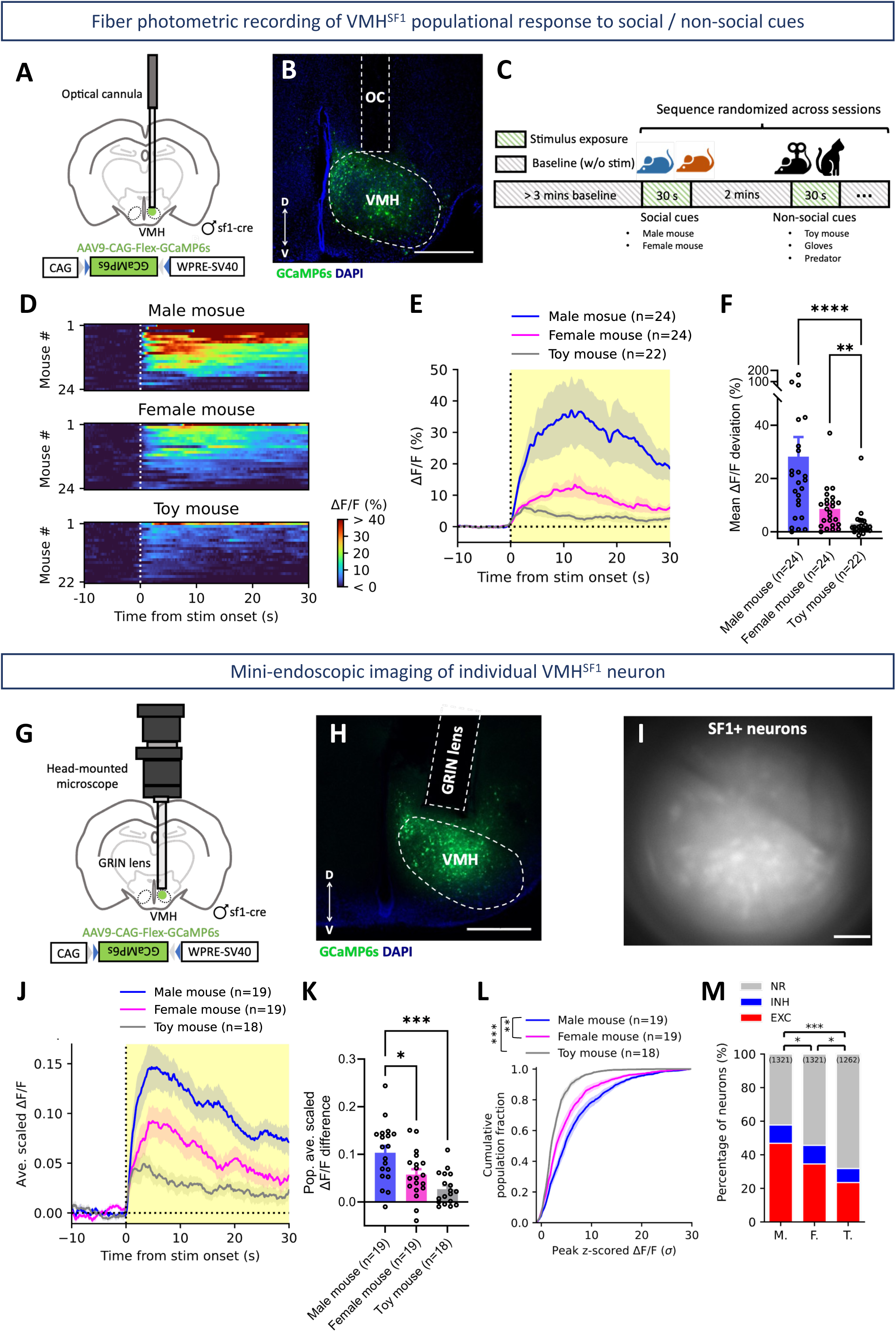
VMH^SF1^ neural population is robustly activated by social cue with male-biased conspecific sex preference. **(A-B)** Schematics and histology of VMH^SF1^ fiber photometric recording. Scale bar: 500 *μm* **(C)** Experimental protocol for stimulus exposure test. **(D)** Heatmaps showing the stimulus-triggered VMH^SF1^ populational Ca^2+^ dynamics. **(E)** Bulk Ca^2+^ responses of VMH^SF1^ neurons evoked by stimulus presentation. **(F)** Averaged stimulus-triggered VMH^SF1^ populational responses. **(G-H)** Schematics and histology for microendoscopic imaging of VMH^SF1^ neurons. Scale bar: 500 *μm* **(I)** Microendoscop field-of-view of a representative mouse VMH. Scale bar: 100 *μm* **(J)** Population Ca^2+^ responses of VMH^SF1^ neurons evoked by stimulus presentations. **(K)** Averaged and scaled stimulus-triggered VMH^SF1^ populational responses. **(L)** Cumulative distribution of neural response amplitudes (Kolmogorov-Smirnov test). **(M)** Fraction of neurons classified as stimulus excited (EXC), inhibited (INH) or non-responsive (NR) (Chi-squared test. M: Male mouse, F: Female mouse, T: Toy mouse).

### Discrepant external stimuli recruit distinct subsets of VMH^SF1^ neurons

To further decipher the conspecific sex-encoding pattern of individual VMH^SF1^ neurons, we inspected how individual neurons responded to distinct conspecific sexes. We found that VMH^SF1^neurons exhibit a wide range of conspecific sex preferences, with 22.1% of neurons exclusively excited by male conspecific and 10.2% exclusively excited by female conspecific (Figure 2A). Moreover, choice probability analysis revealed that 76.7% of VMH^SF1^ neurons showed substantial sex-bias (Figure 2D). This implies that different sexes recruit separable ensembles of VMH^SF1^ neurons and conspecific cues are encoded with sex-specificity, with male-preferring outnumbering female-preferring neurons (Figure 2B-D).

**Figure 2.**
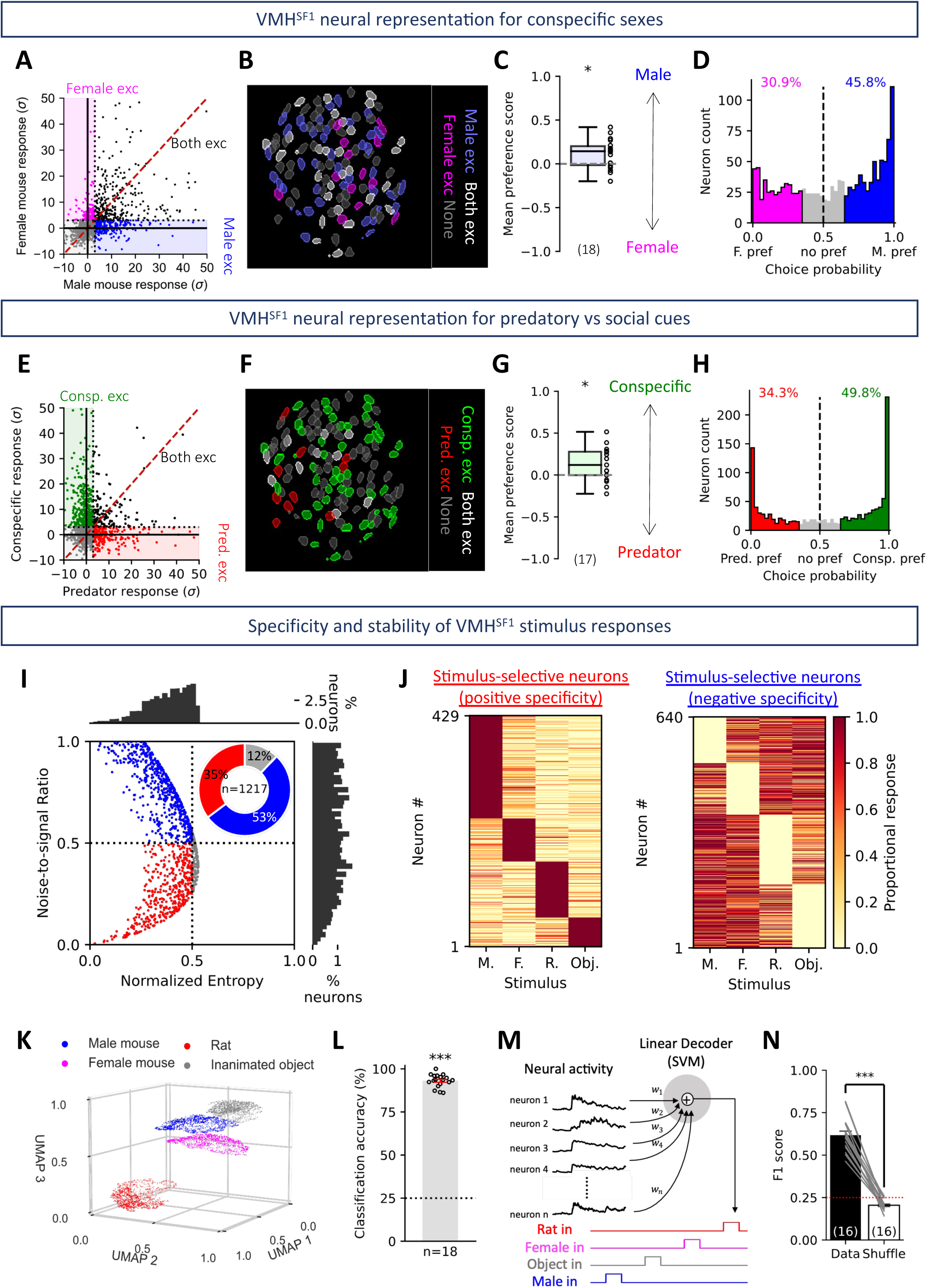
Discrepant external stimuli recruit distinct subsets of VMH^SF1^ neurons. **(A)** Scatter plot showing response amplitude of individual neurons to distinct conspecific sexes (marked color reflects the preference class). **(B)** ROIs from a representative mouse showing spatial distribution of VMH^SF1^ neurons with sex-specific responsiveness. **(C)** Population-averaged conspecific sex preference scores (One-sample Student’s t-test, hypothesized population mean = 0.5). **(D)** Choice probability histogram and percentages of conspecific sex-tuned neurons. (n = 1321 neurons from 18 mice) **(E-G)** Same as **A-C,** but for neuronal response to the social (Pooled from both male and female mice) and predatory (Rat) stimuli. **(H)** Choice probability histogram and percentages of “species-tuned” neurons. (n = 1214 neurons from 17 mice). **(I)** Scatter plot and histograms of neural entropy and noise-to-signal ratio, with the fractions of neurons showing positive (red) or negative (blue) stimulus specificitysummarized in the top-right panel. **(J)** Heatmaps showing the proportional responses of stimulus-selective neurons with (left) positive or (right) negative stimulus specificity. **(K)** UMAP embedding of population responses, colored by stimulus identity. **(L)** Accuracy of KMeans-predicted classification in matching real data class (One-sample Student’s t-test, hypothesized population mean = 25). **(M)** Linear decoder training framework for predicting stimulus identity from population Ca^2+^ dynamics. **(N)** Performance of the decoder at decoding stimulus type from population neural activity.

Studies have shown that predatory stimuli activate VMH^SF1^ neurons^16,37,66^. Nevertheless, whether conspecific and predator recruit unique subsets of VMH^SF1^ neurons in freely-moving mice remains unclear. Hence, we performed the same pair-wise preference assessment for social versus predatory responses. Our results showed that VMH^SF1^ neurons encoded predator and conspecific with a stronger bias: virtually 50% of neurons were exclusively excited by either conspecific or predator (Figure 2E), and more than 80% exhibiting significant preference for conspecific or predator (Figure 2G-H). Therefore, the social-tuned VMH^SF1^ neurons likely form a subgroup distinct from the predator-responsive ensemble (Figure 2F).

We have demonstrated that VMH^SF1^ neurons selectively encode external cues. However, the level of specificity across multiple stimulus categories is unresolvable through pairwise comparison. Hence, we accessed each neurons’ response across four representative stimuli and calibrated the normalized entropy and noise-to-signal ratio correspondingly. Strikingly, the distribution of normalized entropies concentrated far away from 1.0 (population average: 0.3805±0.003, maximum: 0.5304), indicating that VMH^SF1^ neurons exhibited certain level of disparity in response magnitude across different stimuli. Moreover, the noise-to-signal ratio distributed uniformly from 0 to 1, with a population average of 0.5399±0.007 (Fig 2I). Altogether, more than 85% of neurons exhibited positive/negative specificity (Fig 2I), thus manifesting that VMH^SF1^ neurons form stimulus-specific representation for multiple external cues through selective recruitment/exclusion of corresponding neural subsets (Fig 2K, S2A-B). Moreover, as individual neurons showed highly discrepant activity dynamics, stimulus-specific populational neural state could emerge at population level. Hence, we embedded the stimulus-evoked Ca^2+^ dynamics of each neuron and projected them onto a three-dimensional UMAP subspace (Figure 2K). We notice that embedded single-neural responses formed 4 distinct clusters. The stimulus-associated clusters were highly separatable, as group labels generated by KMeans clustering achieved >90% match with true labels (Figure 2L). Therefore, distinct external cues not only recruit discernable subsets of VMH^SF1^ neurons but elicit unique patterns of Ca^2+^ dynamics in each neuron.

To verify the consistency of such stimulus representation, we cross-registered the Ca^2+^ traces of the same VMH^SF1^ neurons on different days. We found that same stimulus elicited similar Ca^2+^ dynamics in most VMH^SF1^ neurons across days (Figure S1A-C). For all representative stimuli, substantial fraction of neurons maintained their response class over time (> 60% across 2 sessions, > 50% across 3 sessions, Figure S1D-E). Despite the overall stable stimulus encoding (Figure S1F), 25% to 34% of neurons responded differently to the same stimulus between sessions (Figure S1G). However, most of the alterations were either switching between responsive and non-responsive, or varying the magnitudes. Furthermore, neurons that showed reversed responses (EXC to INH, INH to EXC) across sessions accounted for less than 3% of neurons with inconsistent response type. To test if the consistent stimulus representation contributes to the decodability of stimulus identities, we employed decoder analysis(Figure 2M). The decoder trained with true stimulus labels successfully annotated all four stimuli, reaching an average F1 score of 0.618±0.024, which was significantly higher than the decoder trained with shuffled stimulus-labels (Figure 2N, S2C-D), showing that VMH^SF1^ neurons encode stimulus identity with chronological stability.

### Pheromonal signal is critical for male-biased conspecific sex response of VMH^SF1^ neurons

With sex-biased conspecific response observed, we next attended to the sensory modality and context required for male mouse presentation to induce robust VMH^SF1^ population response. We introduced an anesthetized male mouse to eliminate social behavioral feedback. Alternatively, we introduced an awake, dangled male mouse to reduce olfactory sensory inputs, especially non-volatile pheromone signals that requires physical interactions (Figure S3A). Replacing an awake male mouse with an anesthetized male mouse triggered comparable response (Figure S3B-C). However, presenting a dangled male mouse significantly reduced the strength of population response (Figure S3B, D), suggesting that the sensory cues released from the male mouse, but not social feedback, are required to fully activate the VMH^SF1^ neurons.

Rodents rely heavily on olfactory cues to interact with conspecifics^20^. The mouse chemosensing system includes the main olfactory system that senses volatile chemosignals and a vomeronasal system specialized for detecting non-volatile signals during direct contact^5,9^. A dangled mouse presented at a distance presumably abolished non-volatile pheromonal signal sensations due to the absence of physical contact, with a relatively mild impact on the strength of volatile olfactory inputs.

To verify the chemosensory modality required for VMH^SF1^ population to respond preferentially to male conspecific, we first removed the chemosensory input from the major olfactory system by selectively ablating major olfactory epithelium (MOE) sensory neurons through chemical treatment (Figure S3E). We found no difference in population sex bias after the treatment (Figure S3F-H). We next abolished male-specific pheromone cues by introducing a castrated male mouse (CM) (Figure S3E) and compared the triggered responses with those evoked by a gonad-intact (G.I) male mouse. CM elicited a weaker population response (Figure S3I-K), alluding that non-volatile pheromones are vital for male-biased VMH^SF1^ conspecific response. To further decipher how pheromonal signals affect conspecific sex representation of each VMH^SF1^ neurons, we introduced gonad-intact/gonadectomized conspecifics of both sexes to test mouse with head-mounted microendoscope (Figure 3A). Without pheromonal cues from conspecifics of both sexes, most of the VMH^SF1^ neurons showed sex-invariant conspecific responses (Figure 3B-D), as a large fraction of sex-specific neurons were turned unresponsive to both sexes when encountering gonadectomized mice (Figure 3E). Moreover, neurons that showed sex-specific activation by CM or OVXF came from a wide variety of sources: 30% retained their preferred sex when facing GI mice, 19.7% reversed their sex preference, and the rest of them turned from sex-invariant (Co-active or Not active) to sex-selective (Figure 3E). The results implied that the absence of pheromonal input altered the sex preference of each neuron differently. Nevertheless, at the population level, the preference for male over female nearly diminished with CM encountered (Figure 3F-G), whereas the absence of putative female-specific pheromones caused no significant effect on population male preference (Figure 3H). We next turned to the sex differences in the embedded neural structure associated with conspecific-elicited Ca^2+^ dynamics (Figure 3I). Facing gonad-intact conspecifics of distinct sexes, projections of stimulus-triggered responses from individual neurons formed visually discernable clusters, while projections associated with different sexes of gonadectomized conspecifics showed significantly left-shifted and overlapped populations (Figure 3J), alluding poor cluster separability. However, binary decoder reached comparable performance in discriminating sex from neural activities even without pheromone cues (Figure 3K). Therefore, pheromonal signals are critical for VMH^SF1^ neurons to establish sex-biased conspecific representation, but not sex discrimination.

**Figure 3.**
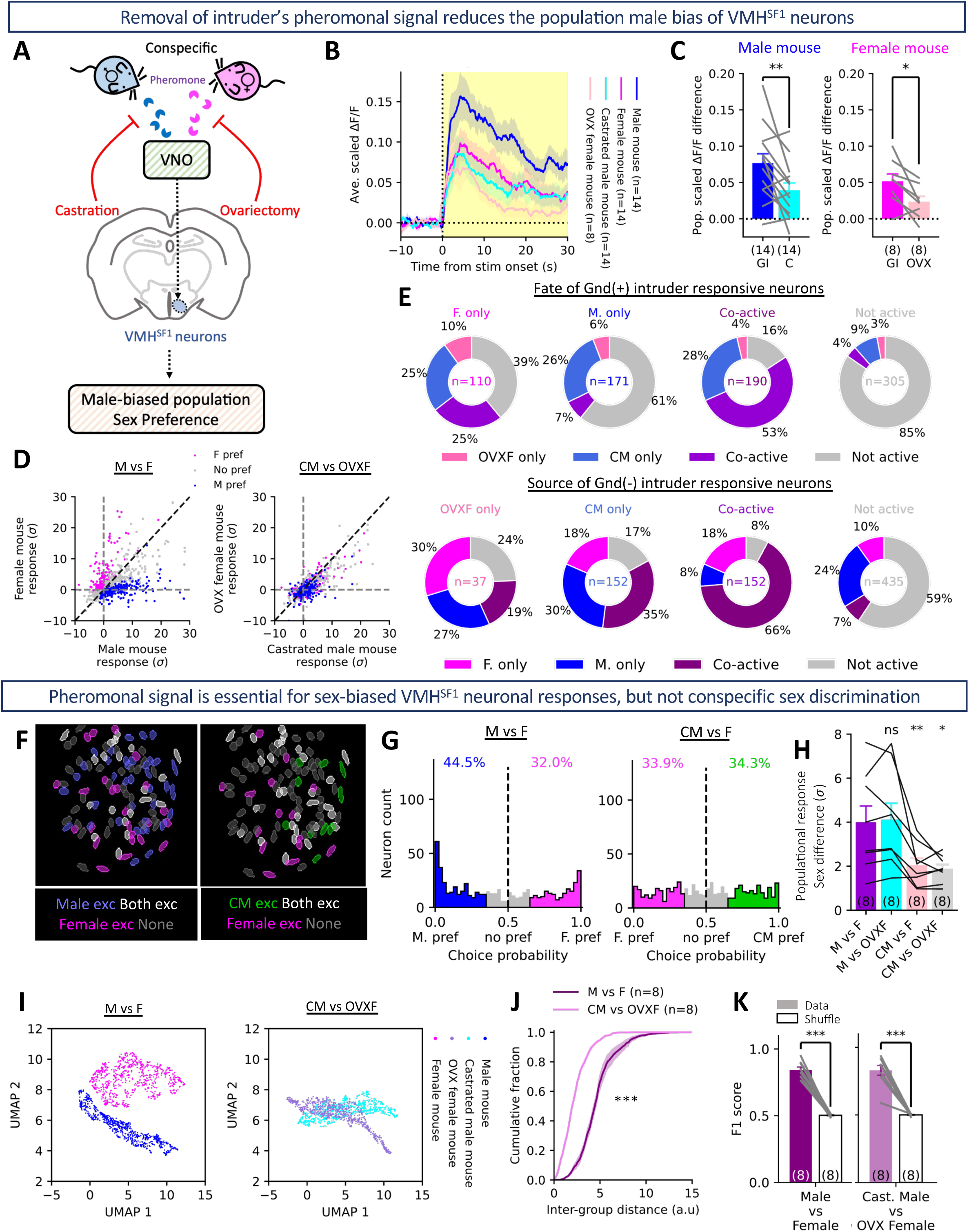
Pheromonal signal is critical for male-biased conspecific sex response of VMH^SF1^ neurons. **(A)** Experimental designs for removing conspecific-emitted pheromones. **(B)** Conspecific-triggered population Ca^2+^ responses of VMH^SF1^ neurons. **(C)** Averaged scaled population response triggered by male (left) or female (right) mouse. **(D)** Scatter plot showing response amplitudes of individual neurons to male versus female mouse (left), or to CM versus OVXF mouse (right). Male-, female-preferring neurons were determined from the left panel and used for marking data in both panels. **(E)** Donut charts showing fates (top) and sources (bottom) of neurons of various sex preferences with/without pheromonal social cues. **(F)** ROIs from a representative mouse showing spatial distribution of VMH^SF1^ neurons with sex-specific or -invariant response to gonad-intact female versus gonad-intact (left) or castrated (right) male mouse. **(G)** Choice probability histogram and percentages of conspecific sex-tuned neurons when encountering gonad-intact female versus gonad-intact (left) or castrated (right) male mouse (n = 776 neurons from 8 mice). **(H)** Individual-averaged sex difference in conspecific-elicited neural responses across various combinations of encountered mice (Paired Student’s t-test with “M vs F” as reference). **(I)** UMAP embedding of population responses to male versus female (left) or CM versus OVXF mouse (right). (n = 776 neurons from 8 mice) **(J)** Cumulative distribution of pairwise distances between male- and female-evoked population responses in the UMAP space. (Kolmogorov-Smirnov test). **(K-L)** Performance of binary decoder in decoding conspecific sexes.

### BNST-VMH pathway is required for male-biased population sex preference of VMH^SF1^ neurons

Our chemosensory ablation experiments suggested that non-volatile pheromonal signals are the key components for VMH^SF1^ neurons to establish male-biased conspecific sex representation. Therefore, VMH^SF1^ neurons likely receive circuit inputs conveying pheromonal information from vomeronasal organ (VNO). Among them, BNST is a vital structure for relaying non-volatile chemosensory information to the medial hypothalamus^8^. During social interaction, BNST is required for sex-recognition and sex-specific innate social behaviors^3^. Moreover, BNST activity is essential for establishment of male-biased conspecific sex representation among VMHvl^ESR1^ neurons^76^.

Anatomically, BNST axons innervate VMH unequally, VMHvl receives intensive innervation, yet the projection targeting VMHdm are relatively sparse^24,75^. However, BNST sends abundant inputs to interneurons throughout VMH shell, a prominent source of local inhibition to VMH core neurons^75^. In addition, studies performing retrograde tracing reported the presence of BNST direct innervations targeting VMH^SF1^ neurons^58,77^. Therefore, VMH^SF1^ neurons likely receive pheromone-related social information directly or indirectly from BNST.

To verify if BNST inputs functionally modulate VMH^SF1^ neurons, we ipsilaterally injected AAV carrying ChrimsonR, a red-shifted excitatory opsin, into BNST and imaged the VMH^SF1^ neurons (Figure 4A and B). An imaging session of empty home cage exploration was employed with multiple sequences of red-light delivery (Figure 4C). Stimulating BNST-VMH pathway at 5 Hz silenced around 5% of the VMH^SF1^ neurons (Figure 4D-E). Strikingly, pathway stimulation at 20 Hz evoked robust time-lock silencing in 61% of VMH^SF1^ neurons, resulting in a strong suppression of population activity (Figure 4D-E), suggesting that BNST-VMH pathway exerts net inhibitory modulation on VMH^SF1^ neurons, functionally confirming the presence of input.

**Figure 4.**
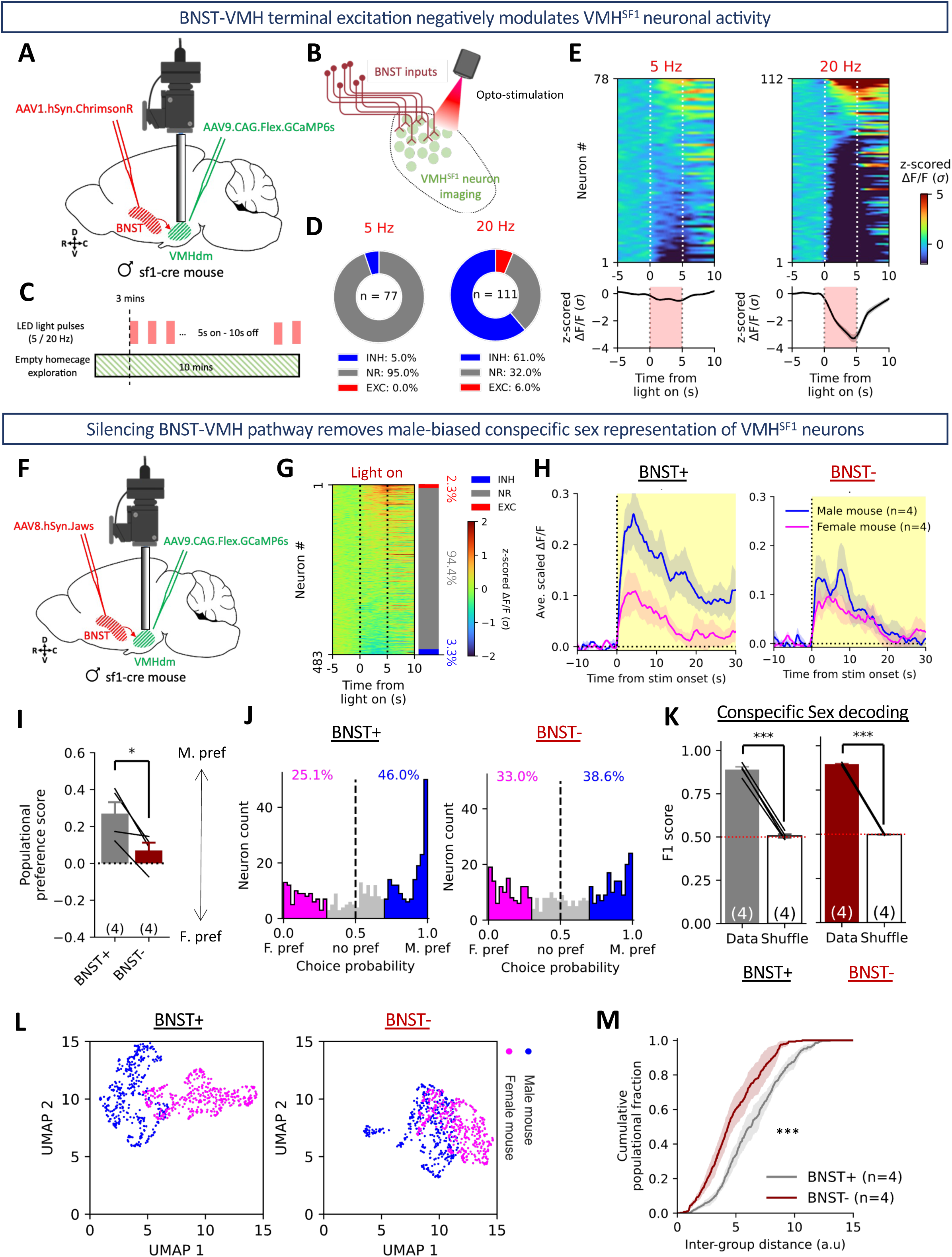
BNST-VMH pathway is required for male-biased population sex preference of VMH^SF1^ neurons. **(A)** A scheme for VMH^SF1^ Ca^2+^ imaging and optogenetic stimulation of BNST-VMH pathway. **(B)** The desired outcome achieved through pathway modulation. **(C)** Experiment design for accessing light-triggered response in VMH^SF1^ neurons,. **(D)** Fraction of VMH^SF1^ neurons showing significant response to BNST-VMH pathway activation at 5 Hz (left) or 20 Hz (right). **(E)** Heatmap (top) and population-averaged Ca^2+^ dynamics (bottom) reflecting the acute response of VMH^SF1^ neurons evoked by 5 Hz (left) or 20 Hz (right) BNST-VMH pathway activation. **(F)** A scheme for VMH^SF1^ Ca^2+^ imaging and optogenetic inhibition of BNST-VMH pathway. **(G)** Heatmap showing acute Ca^2+^ dynamics of VMH^SF1^ neurons evoked by BNST-VMH pathway inhibition, and the fraction of neurons showing significant responses (right). **(H)** Population Ca^2+^ responses of VMH^SF1^ neurons during presentations of male or female mouse, with BNST-VMH pathway functionally intact (left) or optogenetically silenced (right). **(I)** Populational conspecific sex preference with/without intact BNST inputs. **(J)** Choice probability histogram and percentages of conspecific sex-tuned VMH^SF1^ neurons with BNST-VMH pathway functionally intact (left) or optogenetically silenced (right). (n = 483 neurons from 4 mice). **(K)** Performance of decoder in decoding conspecific sexes with (left) or without (right) BNST-VMH input. **(L)** UMAP embedding of population responses to different sexes with (left) or without (right) functionally intact BNST-VMH pathway (n = 483 neurons from 4 mice) **(M)** Cumulative distribution of pairwise distances between male- and female-evoked population responses in the UMAP space. (Kolmogorov-Smirnov test).

Next, to characterize how BNST input influences sex coding of VMH^SF1^ neurons, we adopted loss-of-function approach, excluding the involvement of collateral pathways upon stimulation. Similarly, BNST was transfected with red-shifted inhibitory opsin (Jaws), and VMH^SF1^ neurons were targeted for imaging (Figure 4F). We found that optogenetic silencing of BNST-VMH axon terminals merely modulated 5.6% of imaged neurons (Figure 4G), indicating that BNST-VMH pathway remains quiescent in the absence of salient external stimulus. Subsequently, we performed two sessions of stimulus exposure tests: One without light delivery (BNST+), and the other with red light delivered during stimulus presentation. We found that silencing BNST-VMH pathway during social encounter phenocopied the results acquired when pheromonal inputs were absent: When BNST-VMH pathway was optogenetically silenced, VMH^SF1^ neurons responded to male and female mouse with comparable strength (Figure 4H), and the population sex preference toward male mouse was abolished (Figure 4I-J), with the neuronal sex discrimination remained intact (Figure 4K). Hence, the pheromonal signals likely shape the population sex preference of VMH^SF1^ neurons through BNST-VMH pathway in a direct or indirect manner.

### VMH^SF1^ neurons are strongly tuned to investigation with sex-specificity under social context

With the encoding mechanisms for social cues revealed. We in turn aimed at characterizing the social behavior encoding features. We performed Ca^2+^ imaging during reciprocal social interaction (RSI) test, with behavioral videos simultaneously captured, and social behaviors annotated *post-hoc* (Figure 5A and S6A). We noticed that bulk Ca^2+^ spikes of VMH^SF1^ population were associated with the onset of social-investigation events (Figure S6B-C); social-investigation-associated activations could also be seen in individual neurons (Figure 5B). Furthermore, a significant positive correlation exists between population response magnitude to conspecific upon initial contact, and the percentage of time animal spend investigating throughout the entire interaction period (Figure 5C). To determine the social behavioral coding of VMH^SF1^ neurons thoroughly, we trained a linear SVM decoder for several social behaviors (Figure 5D). We found that activities of VMH^SF1^ neurons encoded multiple social behaviors, yet the behavior decoder trained on investigation achieved vastly better performance than those trained on other social behaviors (Figure 5E-F).

**Figure 5.**
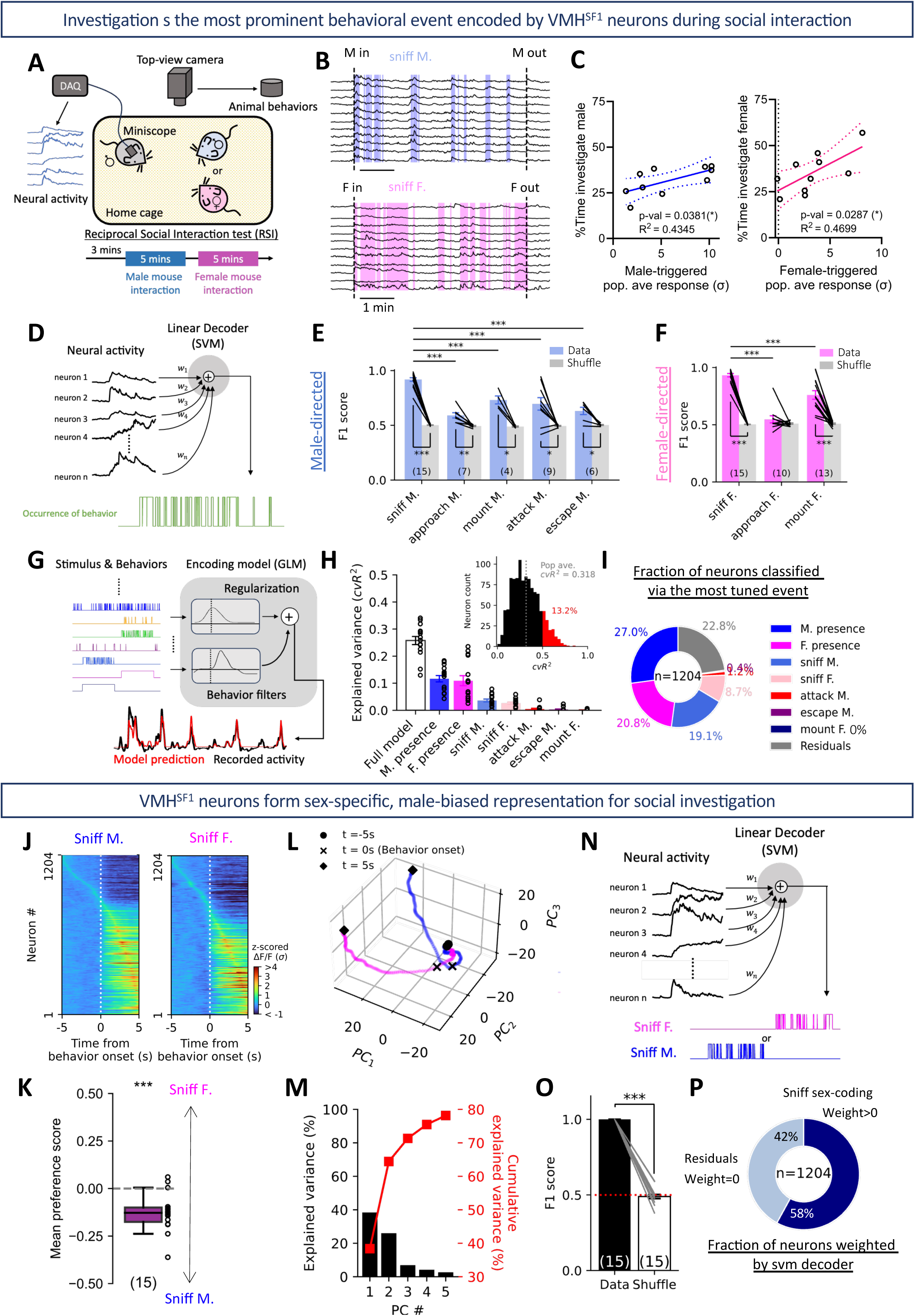
VMH^SF1^ neurons are strongly tuned to social investigation with sex-specificity. **(A)** Experimental procedure and setup for VMH^SF1^ imaging during RSI test. **(B)** Ca^2+^ traces of VMH^SF1^ neurons tuned to male- (top) or female-directed (bottom) investigation. **(C)** Percentage of time spent investigating male/female mouse plotted against male/female-triggered population response upon initial contact. **(D)** Linear decoder architecture for predicting occurrence of specified behavior, trained on neural activity. **(E-F)** Performance of SVM decoder in predicting male- (E) or female-directed (F) social behaviors (Above: Unpaired Student’s t-test with sniff M/F as reference; Beneath: Paired Student’s t-test). **(G)** GLM architecture for detecting neurons selectively tuned to each social behavior/stimulus. **(H)** Distribution of GLM fitting performance (cross-validated R^2^, Top-right) and population-averaged variance explained by full encoding model or simple regression models. (n = 15 mice). **(I)** Fraction of neurons significantly modulated by each regressor. **(J)** Heatmaps of neural Ca^2+^ dynamics associated with male- (left) or female-directed (right) investigation. **(K)** Population-averaged male versus female investigation preference scores (One-sample Student’s t-test, hypothesized population mean = 0.5). **(L)** Investigation-associated population neural trajectories in PC-spanned subspace. **(M)** Explained variance of the first 5 PCs for population response to male/female investigation. **(N)** Linear decoder architecture for predicting male/female investigation. **(O)** Performance of decoder in predicting sniff male/female. **(P)** Fraction of neurons weighted by the decoder. Neurons with non-zero SVM weight were termed the “sniff sex-coding population”.

To further address the behavioral tuning profile of individual neurons, we utilized a generalized linear model (GLM) to characterize the selectivity of each neuron to a variety of stimuli or behavioral events (Figure 5G). The GLM full model could fit 13.2% of neurons with cvR^2^ > 0.5, with a population average of 0.318 (Figure 5H). We further implemented several simple linear regressions to access the performance of single stimulus or behavior in explaining the activity variance of neurons. The GLM full model was able to achieve cvR^2^ of 0.258±0.015, which was significantly higher than all simple regression models. Among all the regressors, conspecific presence contributed the most weight to GLM model fit, with itself fitting the neurons with substantially better performance than behavior regressors (Figure 5H-I). Additionally, male- and female-directed investigation were the behavioral events that contribute the most to GLM fitting, and the major behavioral events individual VMH^SF1^ neurons were tuned to (Figure 5H-I). Hence, our results revealed that, at single-cell level, VMH^SF1^ neural activity variance was explained primarily by conspecific existence, and to a less extend, by social investigation.

Albeit averaged from dozens of events spanned across minutes, VMH^SF1^ neurons showed precise alignment between the onset of activity deviation and social investigation (Figure 5J), signifying a consistent and accurate VMH^SF1^ neural response triggered upon investigation. Moreover, we observed a male-biased population preference in investigation-associated response (Figure 5J-K), resembling passive stimulus representation (Figure 2C-D). To further decipher if investigating conspecifics of different sexes induces discernable patterns of population dynamics, we exploited principal component analysis (PCA) to transform high-dimensional neural data into principal components (PCs) reflecting weighted combinations of all neural activities. We preserved the first 3 PCs (account for 71% of total variance, Figure 5M) for visualizing investigation-associated population dynamics. The PC trajectories associated with male- and female-investigation started separating before the onset of behavior, with the divergence gradually magnified afterward (Figure 5L). By training a linear SVM on single-neural activity, we were able to predict the sex of the investigating conspecific with high accuracy (Figure 5N-O), with 58% of imaged neurons weighted on the sniff male versus sniff female decoder (Figure 5P). Because a smaller fraction of neurons (27.8%) showed selective tuning to male- or female-sniffing, with most neurons (47.8%) merely tuned to the presence of a mouse (Figure 5I); our results imply that VMH^SF1^ neurons achieve sex-specific coding of investigative behavior at the population level.

### Pheromonal and BNST inputs are dispensable for sex-specific investigation coding

Pheromonal signals and functionally intact BNST-VMH pathway were revealed to be essential for maintaining typical pattern of conspecific sex representation in VMH^SF1^ population. Whether such influence remains for encoding investigating different sexes was unresolved. Hence, we carried out imaging in a modified RSI test, in which CM and OVXF mice were introduced instead (Figure S4A). We noticed an apparent reversal in the sex preference of investigation coding. The population response to investigation exhibited clear bias toward OVXF over CM (Figure S4B-D). However, regression analysis revealed that fraction of neurons tuned to investigation targeting CM or OVXF remained consistent (M vs F: 19.1% vs 8.7%; CM vs OVXF: 19.1% vs 8.6%), despite a reversal in the composition of neurons tuned to distinct conspecific sexes (M vs F: 27.0% vs 20.8%; CM vs OVXF: 13.6% vs 34.2%) and a substantially elevated population variance explained by OVXF presence (Figure 5H-I, S4E-F). RSI without conspecific-emitted pheromones also preserved the divergent population dynamics associated with investigating gonadectomized mice of different sexes (Figure S4G-H), and the high decodability of investigating sex from neural activity (Figure S4I). Similar results were obtained when we silenced BNST-VMH pathway during RSI test (Figure S5A). BNST-VMH pathway inhibition reduced the sex bias of investigation-associated activity (Figure S5B-C) and rendered the neurons more tuned to female presence (Figure S5D-E). Nevertheless, the fraction of investigation-tuning neurons, and the ability of neural population to form decodable representation for male or female investigation, were kept unchanged (Figure S5E-F). Hence, we suggest that sex-specific stimulus and behavioral coding of VMH^SF1^ neurons involve discrepant sensory and circuit mechanisms.

### VMH^SF1^ neurons are silenced during consummatory and defensive social behaviors

With social-investigation-encoding features of VMH^SF1^ neurons disclosed, an important argument emerges: As investigation inevitably leads to strengthened sensory inputs, sniffing-associated activity of VMH^SF1^ neurons likely reflects integration of sensory information, rather than a behavior state that facilitates social investigation. Therefore, we explored other social behaviors associated with physical contact, such as aggression or mating. Interestingly, bulk Ca^2+^ activity exhibited substantial escalation upon appetitive but remained silent for consummatory social behaviors (Figure S6D-E). Moreover, conspecific-triggered VMH^SF1^ population response appeared more persistent and intensive if test animal expressed only investigative behaviors throughout the entire session (Figure 6A), whereas population neural activity started declining as aggressive behaviors initiated (Figure 6B). We next inspected the Ca^2+^ dynamics of individual neurons associated with male-directed mount, attack, or female-directed mount. In contrast to net-excitatory response upon social investigation, consummatory behaviors triggered net-inhibitory population response (Figure S6F, H, I). Likewise, both mounting and attacking a male mouse were associated with declining population Ca^2+^ dynamics (Figure 6D, S6J). These dichotomies created a strong population bias toward male investigation over consummation (Figure 6E-G, S6K-L). A similar yet subtle profile of inhibitory population response to consummatory behaviors was seen in female-directed interactions (Figure S6M-O). Although receiving similar level of sensory inputs, VMH^SF1^ neurons were active merely during social investigation, while silenced upon social consummations, suggesting that VMH^SF1^ neurons are recruited specifically in the appetitive phase of social interaction, in which investigation frequently occurs.

**Figure 6.**
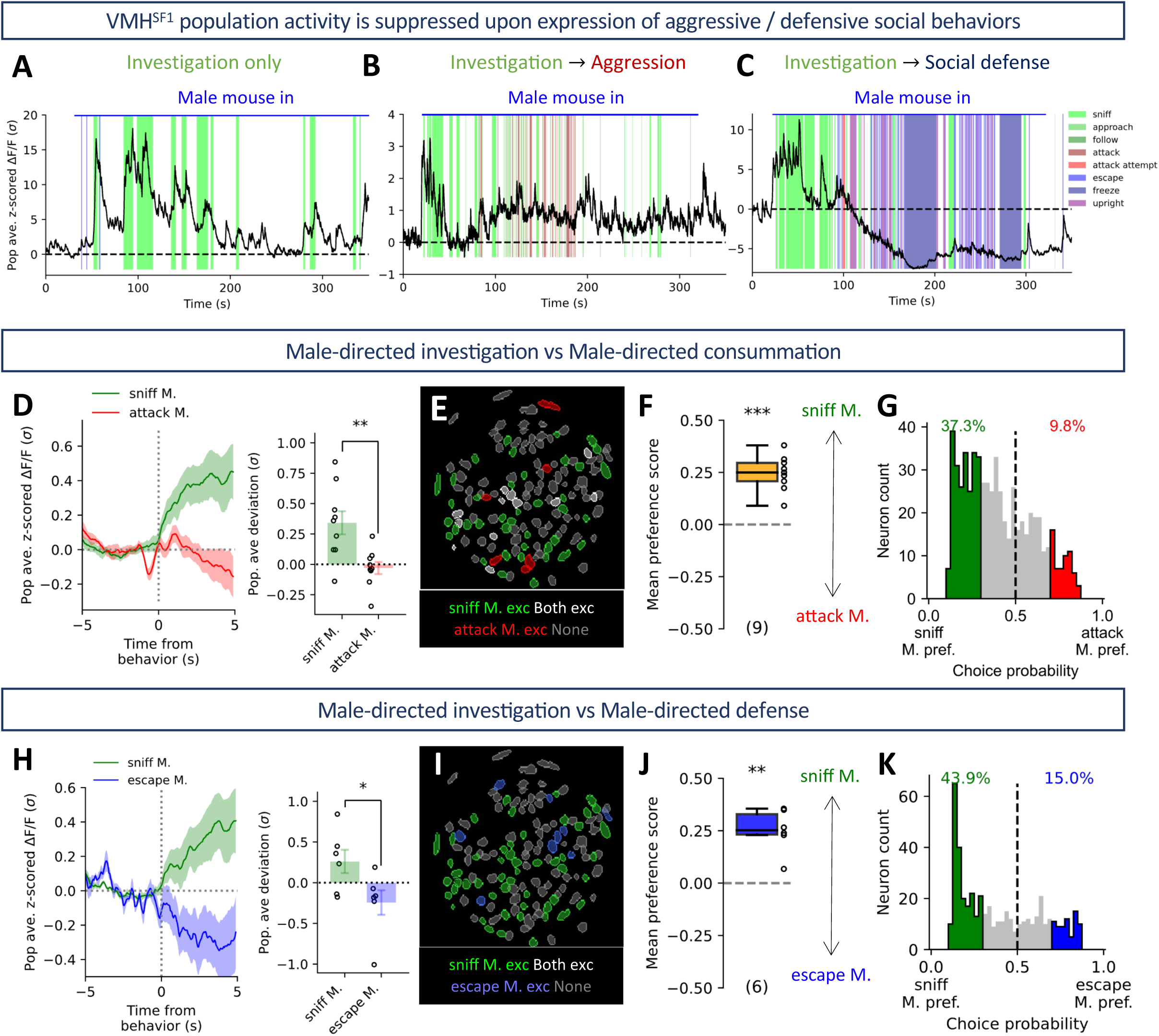
VMH^SF1^ neurons are silenced during consummatory and defensive social behaviors. **(A-C)** Population-averaged Ca^2+^ trace of neurons from male-male interaction trial in which only investigation was expressed (A), aggressive behaviors occurred following initial investigation (B), or strong social defense triggered by social defeat (C) **(D)** (left) Population-averaged Ca^2+^ trace of neurons associated with male-directed investigation/attack. (right) Population-averaged response to male-directed investigation/attack. (n = 9 mice) **(E)** ROIs from a representative mouse showing spatial distribution of VMH^SF1^ neurons with behavior-specific/invariant response to male-directed investigation/attack. **(F)** Population-averaged male investigation versus attack preference scores (One-sample Student’s t-test, hypothesized population mean = 0.5). **(G)** Choice probability histogram and percentages of neurons tuned to male-directed investigation or attack. (n = 652 neurons from 9 mice) **(H-K)** Same as **D-G**, but for male-directed investigation versus escape.

Previous studies have characterized how VMH encodes social and predator defense independently through neurons in the anterior part of VMHvl and VMHdm, respectively^23,31,42,51,57,60,70^. However, the activity pattern of VMH^SF1^ neurons during social defense was unknown. To ensure the test animal was rigorously attacked and defeated, we introduced an aggressive ICR male mouse. Notably, social defeat and the following expression of defensive behaviors were corresponded to a drastic and prolong silencing of VMH^SF1^ population (Figure 6C). Furthermore, VMH^SF1^ neurons exhibited an intense inhibition upon escaping from the aggressor (Figure 6H, S6G), and a strong population preference to male investigation over escape (Figure 6I-K). The results further supported the proposed phase-specific recruitment of VMH^SF1^ neurons during social interaction, as VMH^SF1^ neurons turned silent under social contexts of which further investigation is not needed.

### Social investigation and predator defense recruit discrepant subsets of VMH^SF1^ neurons

Our imaging results have revealed that VMH^SF1^ neurons were well-tuned to social investigation. These behavioral coding features of VMH^SF1^ neurons suggest potential existence of an investigation-prompting behavior state. However, VMH^SF1^ neurons also encode various aspects of predator-oriented defensive internal states^16,37^. The dual-function characteristic of VMH^SF1^ neural population could be implemented either through context-specific recruitment of functionally distinct neural subsets or tuning the output features of the entire population with mixed selectivity (Figure 7A). We have reported that social and predatory stimulus recruited separatable subpopulations of VMH^SF1^ neurons (Figure 2E-J). Nevertheless, whether the discernability in stimulus representation is preserved at the behavioral level was unclear.

**Figure 7.**
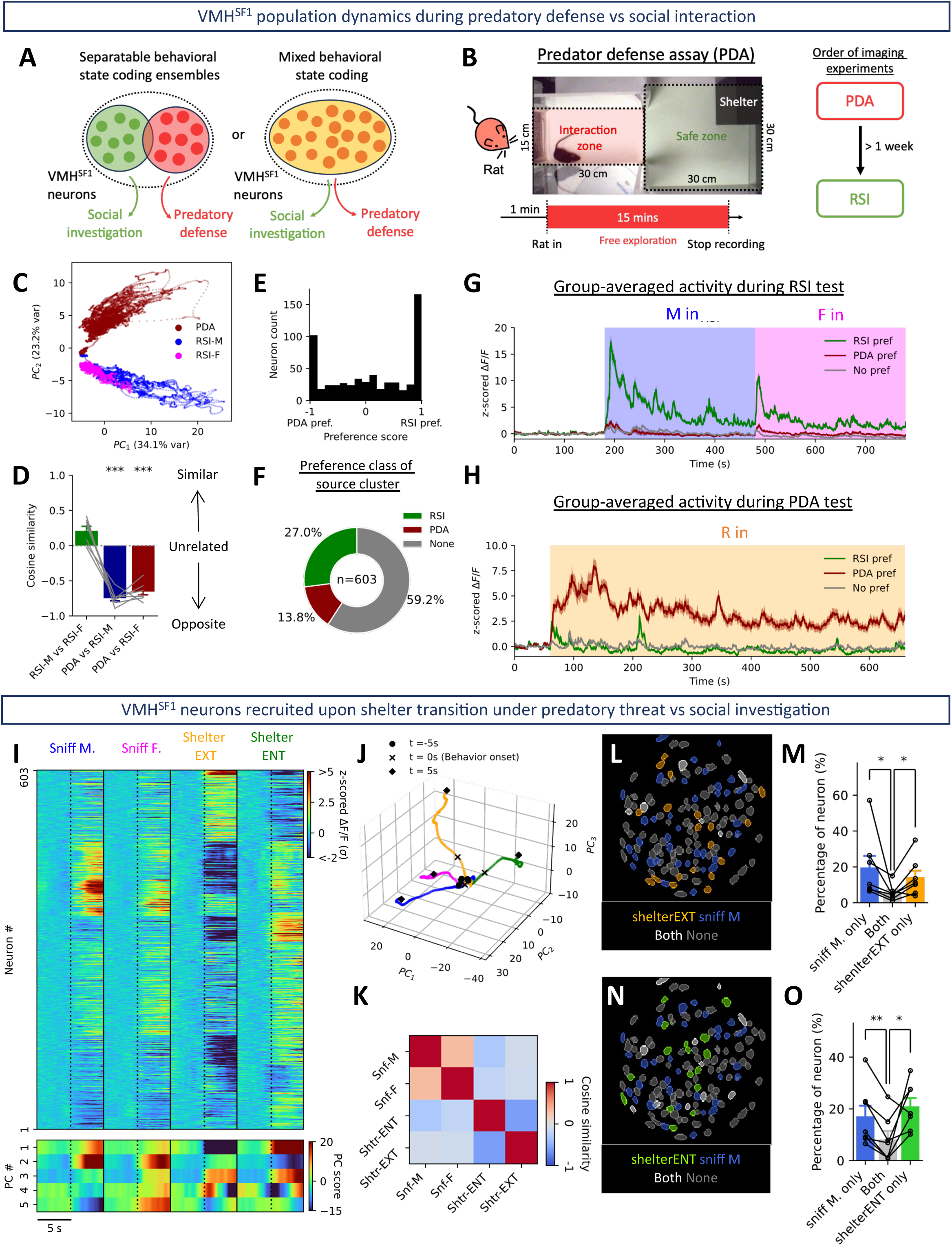
Social investigation and predator defense recruit discrepant subsets of VMH^SF1^ neurons. **(A)** Models for neural population to encode both predator defense and social investigation states. **(B)** Experimental design for the predator defense assay (PDA). **(C)** z-scored Ca^2+^ activities from a representative mouse recorded from RSI and PDA tests projected onto a common PC subspace. **(D)** Cosine similarity between mean trajectory vectors corresponded to different contexts. (n = 7 mice; Paired Student’s t-test with “RSI-M versus RSI-F” as reference). **(E)** Preference score histogram showing the distribution of neuronal trial preference. **(F)** Fraction of neurons with corresponding cluster classified as RSI-, PDA-, or no-pref. **(G-H)** Group-averaged z-score activity recorded from RSI (G) or PDA (H) test of neural subgroups with distinct context preferences defined in **F**. **(I)** Trial-averaged, z-scored Ca^2+^ activities (top) of registered neurons under 4 behavioral conditions, transformed into time-evolving PC scores. (bottom) **(J)** Population neural trajectories under 4 behavioral conditions in the three-dimensional subspace spanned by PC1∼3. **(K)** A heatmap showing the cosine similarity of mean PC trajectory vectors across behavioral conditions. **(L)** ROIs from a representative mouse showing spatial distribution of VMH^SF1^ neurons with distinct responsiveness for shelter exit in PDA or male-directed social investigation. **(M)** Percentage of neurons that are selectively responsive or co-responsive to shelter exit or male-directed social investigation. (n = 7 mice) **(N-O)** Same as **L-M**, but for shelter entry versus male mouse investigation.

Hence, we designed a predator defense assay (PDA), in which test animal was allowed to access a rat or to stay in a shelter away from rat (Figure 7B). From same group of neurons, we co-registered the Ca^2+^ traces imaged in PDA with those captured from RSI test. We employed hierarchical clustering over all neurons based on patterns of merged activity and noticed a visually perceivable context-specificity in multiple neural clusters (Figure S8A). Therefore, we seek to examine if the population Ca^2+^ dynamics from two behavioral tests had discernible features. We performed PCA on the merged population activities of each mouse and divided the transformed neural data into sections corresponded to PDA, male mouse interaction (RSI-M), and female mouse interaction (RSI-F). Despite having similar origins during baseline recording, the full-trial PC trajectory after PDA initiation occupied a unique territory, with minimal spatial overlap with those spanned by neural trajectories of RSI-M and RSI-F (Figure 7C). To test if the divergence in neural states elicited in PDA versus RSI is seen across mice despite variations in behavioral expressions, we calibrated the cosine similarity of neural trajectory mean vectors for pairs of conditions. Cosine similarity between RSI with both sexes and PDA was significantly lower than the that between RSI-M and RSI-F (Figure 7D), demonstrating that under predatory threat, VMH^SF1^ neuron express patterns of population activity vastly different from social context. At the cellular level, some neural clusters were specifically active in either PDA or RSI test (Figure S8A). By calibrating the preference scores utilizing mean activity deviation from test-associated baseline (Figure 7E), 40.8% of neurons were identified as RSI- or PDA-preferring (Figure 7F), which were persistently active with context specificity (Figure 7G-H).

To gain further insight into the behavioral representation under predatory threat, we explored individual neuronal coding of predatory threat (out of shelter/exit shelter) and safety (in shelter/enter shelter). Consistent with recent studies^16^, more neurons were preferentially activated by shelter exit or explorations out of shelter (Figure S7B-C), causing a net populational excitation upon shelter exit, and inhibition upon entry (Figure S7D). Besides shelter transitions, VMH^SF1^ neurons also responded to defensive or risk assessment behaviors with unique dynamics (Figure S7F). Moreover, the responsiveness of VMH^SF1^ population to initial rat encounter tightly reflected the behavioral defensiveness of animal (Figure S7E), altogether suggesting that VMH^SF1^ population activity dynamically tracks the animal’s defensive state.

Subsequently, we paired the Ca^2+^ activities of shelter transition in PDA with those associated with social investigation (Figure 7I). Subsequently, we carried out PCA and preserved the first 3 PCs (explained 63.95% of total variance) for visualizing neural trajectory under four behavioral conditions. We noticed that social investigation and shelter transition under predatory threats corresponded to vastly divergent neural dynamics (Figure 7J). To further determine the extent of separation, we included the first 5 PCs (explained 78.53% of total variance) and calibrated the cosine similarity of mean trajectory vectors for all combinations of behavioral conditions (Figure 7K). Positive cosine similarity between male and female investigation indicated that the two behavior-associated neural trajectories shared a common subspace. Strong negative similarity between shelter exit and entry suggested anti-correlated trajectories occupying distinct manifolds. In contrast, near-zero similarity between social investigation and shelter transitions implied orthogonal manifolds. These results demonstrate that VMHSF1 neurons exhibit structured population dynamics, with social and defensive behaviors engaging distinct neural dimensions.

Independence between neural manifolds could be achieved by recruiting distinguishable groups of neurons^27^. Therefore, we examined if shelter-transition and social investigation draft separable neuron ensembles. Neurons exclusively respond to shelter transition or male mouse investigation significantly outnumbered dual-responsive neurons (Figure 7L-O, S8D, F). Comparable recruitment pattern was seen for female investigation versus shelter transitions (Figure S8B, C, E, G), suggesting that VMH^SF1^ neurons encoding predatory defense or social investigation were distinct subpopulations.

### VMH^SF1^ neurons modulate the level of engagement in social investigation

Our observational results suggested that a subset of VMH^SF1^ neurons is tuned to social stimuli and encodes social investigation. However, whether VMH^SF1^ neurons exert functional influence on social investigation remains elusive. To modulate neural excitability persistently, we performed chemogenetic manipulation on VMH^SF1^ neurons utilizing the DREADD technology^56^: AAV carrying inhibitory DREADD receptor (hM4Di) or fluorescence control (mCherry) was delivered to bilateral VMH of SF1-Cre mouse (Figure 8A). Next, we implemented consecutive trials of RSI tests, each with either CNO or Vehicle i.p injected 60 mins before test initiation (Figure 8B). In comparison to the Vehicle trial, CNO treatment on mice with hM4Di resulted in less engagement in social interaction and reduced investigation time toward conspecifics of both sexes (Figure 8C, D). In addition, with CNO treatment, a reduction in investigation frequency and shortening of each investigation bout were seen during male-male interaction but not male-female interaction. The CNO-induced modulations on social investigation were seen in mice with hM4Di but not mCherry-only control (Figure 8E-H).

**Figure 8.**
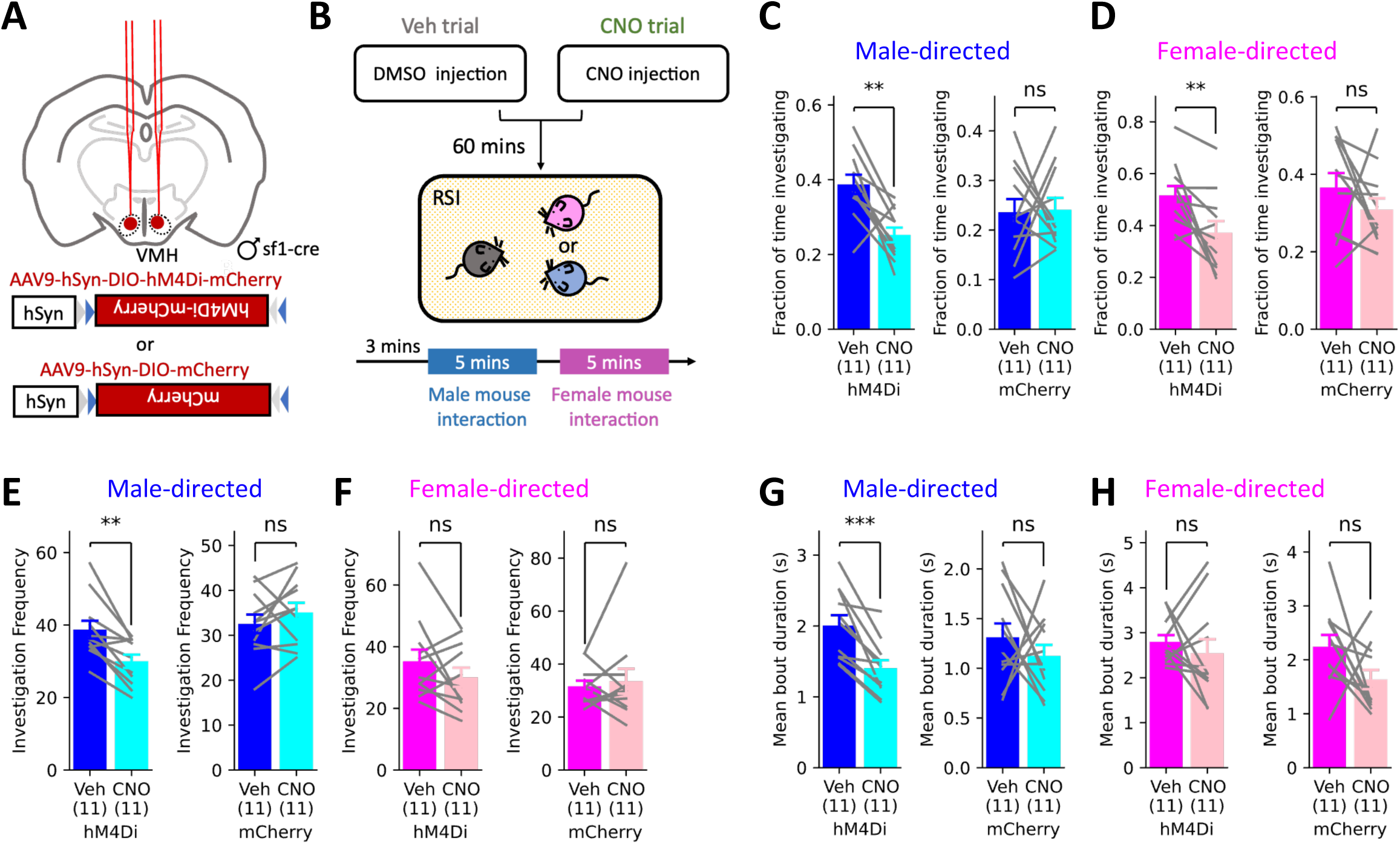
VMH^SF1^ neurons modulate the level of engagement in social investigation. **(A)** A scheme for bilateral SF1-specific transfection of inhibitory DREADD receptor (hM4Di) or fluorescence control (mCherry) in the mouse VMH. **(B)** Experimental design for RSI test with DREADD manipulation targeting VMH^SF1^ neurons. **(C-D)** Fraction of time mice spent investigating a male (C) or female (D) mouse. **(E-H)** Same as **C-D**, but for investigation frequency (E-F) or duration of investigation bout (G-H).

## Discussion

Proper action toward encountered conspecifics requires efficient investigation. Using cell-type-specific monitoring of neuronal Ca^2+^ dynamics in freely behaving mice, we identified a unique subset of VMH^SF1^ neurons that are tuned to social cues with sex specificity and encode social investigation. Silencing the entire population of VMH^SF1^ shortened the duration of social investigation, further supports the functional involvement of VMH^SF1^ neurons in social interaction.

### Sensory and circuit mechanisms underlying VMH^SF1^ neuronal representation of conspecific sex

In line with other studies^11,16,37^, our results showed that VMH^SF1^ neurons encode stimulus identity via recruiting distinct neural subpopulations. Nonetheless, we further revealed that VMH^SF1^ neurons possess comparable specificity in representing different conspecific sexes, with a sturdy male preference resembling VMH^ESR1^ neurons^36,55,76^. Moreover, we identified conspecific-derived, non-volatile chemical signals as a potent sensory input for VMH^SF1^ neurons to exhibit male-favoring population response. Absence of circulating sex hormone due to gonadectomy shall remove most of the sex-specific pheromone production^2,38,65^, and therefore, accessory olfactory circuit downstream of VNO that senses non-volatile pheromones likely involve predominantly in shaping conspecific sex bias of VMH^SF1^ neurons^8,12,73^. However, the influence of volatile chemosensory inputs, though subtle, could not be entirely excluded, as emission of volatile pheromones was also diminished in gonadectomized conspecifics. Additionally, synthesis of some sex-specific chemical signals is independent of circulating sex hormone^25,74^. Hence, genetic disabling of pheromone sensing shall aid accessing the precise contribution of volatile and non-volatile pheromone signals.

Apart from pheromonal sensory cues, intact BNST-VMH pathway activity also helps establish male-biased conspecific response among VMH^SF1^ neurons. Akin to VMH^ESR1^ neurons^76^, VMH^SF1^ neurons are less responsive to male conspecific with BNST inputs silenced. Furthermore, VMH^SF1^ neurons remained capable of decoding conspecific sex without BNST input, alluding that BNST input and pheromones shape the sex-biased neural representation of VMH^SF1^ neurons without hindering sex discrimination. Despite that optogenetic excitation of BNST terminals in VMH causes net inhibitory modulation on a large fraction of VMH^SF1^ neurons, and that BNST neurons possess an intrinsic female-biased conspecific sex response^3,76^, silencing BNST-VMH pathway upon conspecific presentation failed to create an expected disinhibitory effect on female-induced responses among VMH^SF1^ neurons, as magnitudes of female-evoked population response remained (Figure 4). Thus, we speculate the presence of alternative inhibitory inputs as a source of compensation. In addition, BNST is not the only source of chemosensory input to VMH. MeA is capable of processing pheromonal signals and relaying sensory information to VMH^4,33,45,53,73^. Besides, MeA serves as a primary source for VMH^SF1^ neurons to receive processed sensory cues reflecting predator presence^53^. Therefore, how MeA inputs shape sex representation of VMH^SF1^ neurons worths further investigation.

### VMH^SF1^ neurons encode multiple behavior-driving internal states

Imaging studies suggest that VMH^SF1^ neurons encode a predator-specific, persistent internal state that drives innate defensive responses, and a general state reflecting neophobia or arousal^16,37^. In this study, we propose that a conspecific-evoked, investigation-prompting behavioral state is encoded by a subgroup of VMH^SF1^ neurons distinct from the predator-tuned subpopulation driving defensive responses. These social-tuned VMH^SF1^ neurons are persistently, and selectively activated by social external cues, and the strength of their activities reflects the level of engagement in social investigation. The features suggest that the social-tuned VMH^SF1^ neurons encode an behavior-driving internal state distinct from arousal state upon novel item confrontation^1^. Therefore, VMH^SF1^ neurons are not homogeneous, but a supercluster of spatially intermixed neural subclasses, each with its preferred stimulus category and unique behavioral function. Interestingly, studies performing VMH^SF1^ neuron stimulation exclusively resulted in defensive behavioral responses^37,43,69,71^, even though social-tuned VMH^SF1^ neural subsets are bound to be recruited simultaneously. The functional dominance of defense-driving neural subclass aligns with previous studies showing that appetitive behaviors, which make the animal more susceptible to predation, are often actively suppressed by defensive state^43,54,64^. For VMH neurons with mixed populations encoding conflicting behavioral states, studies have shown that the behavioral outcome of optogenetic stimulation is mostly determined by the larger neural subgroup^36,48,76^. As predator cues evoke stronger VMH^SF1^ population response^37^, predator-tuned VMH^SF1^ neurons likely dominate the behavioral effect of optogenetic stimulation. Alternatively, activation of defense-driving VMH^SF1^ neurons could in turn recruit feed-forward inhibition on other VMH^SF1^ neural subsets. As nearly all VMH^SF1^ neurons are glutamatergic^68^, the inhibition is likely mediated by surrounding structures, such as VMH shell, dorsomedial hypothalamus (DMH), tuberal nucleus (TU), or anterior hypothalamus (AHN). Both mechanisms enable the VMH^SF1^ neurons to prioritize defensive over social state in the presence of life-threatening predators. Altogether, our results further disclose the functional diversity of VMH^SF1^ neurons in modulating innate behaviors.

### Functional interplay between distinct VMH neural subclasses

Our results demonstrate that VMH^SF1^ neurons respond to appetitive social investigation in a comparable manner as VMH^ESR1^ neurons, both activated upon conspecific-directed sniffing with male-biased sex preference^30,36,55,76^. Moreover, a neuron subgroup co-expressing SF1 and ESR1 are found in anterior VMH^21^. To minimize the inclusion of VMH^ESR1^ neurons into imaged population, our target coordinate for optical cannula or GRIN lens implantation was set at middle VMH along A-P axis, with *post-hoc* histological confirmation. Furthermore, upon consummatory or defensive social behaviors, our imaged VMH^SF1^ neurons show unique inhibitory dynamics, which are vastly discrepant from VMH^ESR1^ neurons, which are virtually activated^36,50,70,78^. It appears that VMH^SF1^ neurons and VMH^ESR1^ neurons are recruited in distinct phases of social interaction. As social investigation offers sufficient information for terminal decision, other neural substrates, such as VMH^ESR1^ neurons, take over for further behavioral control, switching off investigation-driving modules. On the other hand, weak activation of VMH^SF1^ neurons abrogates ongoing male aggression without causing robust freezing or activity burst^43^, implying that SF1- and ESR1-expressing neurons are able to employ reciprocal inhibition, potentially through feedback inhibition from neighboring regions, under certain behavioral conditions. Moreover, the presence of abundant recurrent synaptic connection and neuropeptide signaling among VMH neurons^37,49,59^ permits bi-directional flow of information between SF1- and ESR1-expressing neurons, which could give rise to equivalent sniff-associated Ca^2+^ dynamics in both populations.

Hence, it will be interesting to further explore the structural, functional, and behavioral features of these proposed intra-nucleus or inter-nucleus circuit to understand the communication between SF1- and ESR1-expressing neurons in the VMH.

## Resource availability

### Lead contact

Further information and request for resources and reagents should be directed to and will be fulfilled by the lead contact, Shi-Bing Yang.

### Materials availability

This study did not generate new unique reagents.

### Data and code availability

All reported data and code used in this paper, as well as additional information required to reanalyze the data, will be shared by the lead contact upon request.

## Acknowledgements

The authors thank the Animal Core, the Pathology Core and the Common Facilities Core Laboratory at the Institute of Biomedical Sciences, Academia Sinica, Taiwan. This work was supported by the Institute of Biomedical Sciences at Academia Sinica, the National Science and Technology Council (111-2320-B-001-008-MY3, 112-2321-B-002-024, and 111-2314-B-001-004 to S.-B.Y.), and the National Health Research Institute (NHRI-EX113-11336SI to S.-B.Y.).

## Author contributions

S-C. L. and S-B. Y. Conceptualized and designed the study. S-C. L. conducted all the experiments, data analysis and figure preparation. T-A. S., H-Z. L. and J-Y. H. contributed to behavioral annotations. S-B. Y supervised the project. S-C. L. and S-B. Y wrote and revised the manuscript.

## Main Figure legends

Unless otherwise noted:

- All Ca^2+^ traces are plotted as Mean ± s.e.m.
- Shaded regions in time-series panels indicate stimulus presentation or behavioral occurrence.
- For linear regression: Dotted lines represent 95% confidence intervals for the regression fits.

## STAR Methods

### Key resources table

| REAGENT OR RESOURCE | SOURCE | IDENTIFIER |
| --- | --- | --- |
| Bacterial and virus strains |  |  |
| AAV9-CAG-Flex-GCaMP6s-WPRE-SV40 | Chen et al.<sup>13</sup> | Addgene viral prep # 100842-AAV9 |
| AAV1-hSyn-ChrimsonR-tdTomato | Klapoetke et al.<sup>40</sup> | Addgene viral prep # 59171-AAV1 |
| AAV8-pAAV-hsyn-Jaws-KGC-GFP-ER2 | Choung et al.<sup>19</sup> | Addgene viral prep # 65014-AAV8 |
| AAV9-hSyn-DIO-hM4D(Gi)-mCherry | Krashes et al.<sup>41</sup> | Addgene viral prep # 44362-AAV9 |
| AAV9-hSyn-DIO-mCherry | Bryan Roth | Addgene viral prep # 50459-AAV9 |
| Chemicals, peptides, and recombinant proteins |  |  |
| Paraformaldehyde (PFA) | Sigma-Aldrich | 16005 |
| Phosphate Buffered Saline (PBS) | PRO Tech | BF203 |
| Clozapine N-oxide | Sigma-Aldrich | SI-C0832 |
| DMSO | Sigma-Aldrich | D-2650 |
| Dichlobenil | Sigma-Aldrich | D57558 |
| Zoletil® 50 | Virbac Taiwan | 05393 |
| Rompun | Bayer Korea Ltd. | CAPROM10m l-L-003 |
| Caprofen | China chemical & pharmaceutical | 3240912 |
| Ampicillin | Y F Chemical Corp. | 4-032-01-2(9) |
| Dexamethasone | China chemical & pharmaceutical | Y20-0025 |
| Isoflurane | Panion & BF Biotech Inc. | DG4900250B |
| Experimental models: Organisms/strains |  |  |
| Mouse: SF1-Cre | Dhillon et al.<sup>22</sup> | RRID: IMSR_JAX:012462 |
| Mouse: C57BL/6J | IBMS, Academia Sinica | N/A |
| Mouse: BALB/c | NLAC | RMRC11001 |
| Mouse: Bltw:CD1(ICR) | IBMS, Academia Sinica | N/A |
| Rat: Bltw:CD(Sprague-Dawley) | BioLASCO Taiwan | N/A |
| Rat: Crl:LE(Long-Evans) | NLAC | RMRC21006 |
| Software and algorithms |  |  |
| GraphPad Prism 9 | GraphPad software | RRID: SCR002798 |
| ezTrack | Pennington et al.<sup>52</sup> | https://github.com/denisecailab/ezTrack |
| BORIS | Friard et al.<sup>29</sup> | https://www.boris.unito.it |
| Machine Vision Software MVS V4.4.0 | HIKROBOT | https://www.hikrobotics.com/en/machinevision/service/download/ |
| Inscopix Data Acquisition Software (IDAS) | Inscopix | https://www.inscopix.com/software-analysis#software_idas |
| Inscopix Data Processing Software (IDPS) | Inscopix | https://www.inscopix.com/software-analysis#software_idps |
| Doric Neuroscience Studio V5 | Doric Lenses | https://neuro.doriclenses.com/products/doric-neuroscience-studio-v5 |
| Axon<sup>TM</sup>pCLAMP<sup>TM</sup> Clampfit 10.7 | Molecular Devices | https://support.moleculardevices.com/s/article/Axon-pCLAMP-10-Electrophysiology-Data-Acquisition-Analysis-Software-Download-Page |
| Python | Python Software Foundation | RRID:SCR_008394 |
| Other |  |  |
| Mounting medium with DAPI | abcam | ab104139 |
| Adhesive glass slide | Matsunami | PRO-11 |
| Adhesive glass slide | Matsunami | APS-14 |
| Vibratome | Campden Instruments | Model 5100mz |
| Flow-It™ ALC | Pentron Clinical | N11B |
| Super-Bond C&B | Sun Medical | K0261 |
| Automated rotational stereotaxic | RWD | 71000-X |
| Heating pad | SUPERTECH | TMP-5b |
| Drill set | FOREDOM | 51449 |
| Nanoliter Injection Pump | RWD | R-480 |
| Glass filament | A-M Systems. Inc. | 615000 |
| Integrated GRIN lens | Inscopix | #100-005620 |
| Fiber optic cannula | inper | FOC-B-1.25-400-0.50-7.0 |
| Mono fiber optic | inper | MFO-F-W2.5-400-0.50-200 |
| Optical fiber patch cord | Doric Lenses | D202-4070 |
| Fiber-optic rotary joints | Doric Lenses | FRJ_1x1_FC-FC |
| Confocal microscope | Zeiss | LSM700 |
| Epifluorescent microscope | Zeiss | Axioscope A1 |
| nVista3.0 | Inscopix | https://inscopix.com/nvista-system/ |
| nVoke2.0 | Inscopix | https://inscopix.com/nvoke-system/ |
| Behavioral camera | HIKROBOT | MV-CS016-10UC |
| Fiber photometry console | Doric Lenses | FPC |
| LED driver | Doric Lenses | LEDD_2 |
| Femtowatt photoreceiver | Newport | Model 2151 |
| 405 nm violet LED | Doric Lenses | CLED_405 |
| 465 nm blue LED | Doric Lenses | CLED_465 |

## Experimental model and subject details Animals

All experimental procedures, with live animal usage included, were approved by the Institutional Animal Care and Use Committee (IACUC) at Academia Sinica (24-08-2258). The SF1-Cre transgenic mice were obtained from the Jackson Laboratory (Stock no: 012462; RRID: IMSR_JAX:012462) and maintained as heterozygotes in the C57BL/6J genetic background. Heterozygous male mice aged between 6 and 40 weeks were used in this study. Since VMH exhibits sexually dimorphic gene expression profile^15,35^, structural and functional characteristics of VMH^SF-1^ neurons might differ between sex and affected by estrous cycle in female mice. Hence, all the animals we used for conducting experiments in this study were male mice. Wild type, white-coat mice under inbred (BALB/c) or outbred (Bltw:CD1(ICR)) genetic background were purchased at the age of 4-6 weeks from the National Laboratory Animal Center (NLAC) or animal facility at the Institute of Biomedical Sciences in Academia Sinica, respectively. These wild type mice were group-housed and used as social stimuli or intruders at the age above 8 weeks. Sprague-Dawley (SD) or Long-Evan (LE) male rats used as predator were purchased from NLAC and BioLASCO Taiwan, respectively at the age of 6-8 weeks. Rats weighted above 600 g or aged above 16 weeks were chosen for behavioral tests. Animals were housed in a SPF (specific pathogen free) facility and maintained under a controlled environment (22.0 ± 2℃, with a 12hr light/dark cycle) with *ad libitum* access to food and water. Mice without surgery treatments were group-housed with up to 5 same-sex littermates per cage, unless otherwise required and explicitly mentioned.

### Viruses

For fiber photometry and microendoscopic Ca^2+^ imaging, SF1-Cre^+/-^ mice were unilaterally injected in VMHdm with 50 nL AAV9-CAG-Flex-GCaMP6s-WPRE-SV40 (Addgene 100842-AAV9, 3.28 × 10^12^ genomic copies / ml). Apart from GCaMP6s delivery to VMHdm, mice for optogenetic BNST-VMH pathway excitation were injected in ipsilateral BNST with 35 nL AAV1-hSyn-ChrimsonR-tdTomato (Addgene 59171-AAV1, 1.0 × 10^13^ genomic copies / ml), whereas mice for optogenetic BNST-VMH pathway inhibition were injected in ipsilateral BNST with 35 nL AAV8 pAAV-hsyn-Jaws-KGC-GFP-ER2 (Addgene 65014-AAV8, 1.0 × 10^13^ genomic copies / ml). For chemogenetic manipulation experiments, SF1-Cre^+/-^ mice were bilaterally injected in VMHdm with 50 nL of 5× diluted AAV9-hSyn-DIO-hM4D(Gi)-mCherry (Addgene 44362-AAV9, 2.50 × 10^13^ genomic copies / ml) or 25 nL of 10× diluted AAV9-hSyn-DIO-mCherry (Addgene 50459-AAV9, 1.20 × 10^14^ genomic copies / ml) as control.

## Method details

### Behavior tests

All behavioral experiments were carried out in the custom-designed arena (Figures 8, S7-S8) or the animals’ home cage (Figures 1-7, S1-S6, S8) during the animal’s subjective light period. Top view videos were filmed at 20 Hz using high speed camera controlled by Machine Vision Software MVS V4.4.0 (HIKROBOT). Before initiating behavioral experiments, test mice were co-housed with sexually naïve, strain-matched, wild type female mouse for a week, followed by single housing for at least 3 days. To minimized experimenter-induced stress during behavioral experiments, mice were tamed by experimenter for at least 3 consecutive days.

### Stimulus exposure test

As illustrated in Figure 1D. Test animal was placed into a cover-removed home cage with fresh bedding. 10 minutes of acclimatization was followed by 3 minutes of baseline recording. Unless specifically mentioned, external stimuli were directly placed into the center of home cage for 30 seconds, allowing test animal to freely interact with. After removal of each stimulus, an interval for at least 2 minute was introduced before presenting the next stimulus. Stimuli delivered could be divided roughly into two categories: Social cues included awake novel mice of different sexes, anesthetized male mouse (Figure S3), castrated male or ovariectomized female mouse (Figure 3, S3-S4), and awake male mouse dangled by experimenter, making physical social interaction unavailable (Figure S3); Non-social cues included inanimate objects such as a toy mouse with equivalent size of a living mouse, or an awake rat confined in a rat cage as predatory stimulus. The sequence of stimuli exposure was randomized across sessions to rule out adaptation or experience-dependent factors in stimulus-triggered neural responses.

### Reciprocal social interaction test (RSI)

Test animal was placed into a cover-removed home cage with fresh bedding. After 10 minutes of acclimatization and 3 minutes of baseline recording, a novel male mouse was introduced, allowing free interaction for 5 minutes before removal. After another 2 minutes, a female mouse was introduced for 5 minutes.

### Predator defense assay (PDA)

To elicit naturalistic and flexible defensive behavioral response in test animals, we designed a customized apparatus. As shown in Figure 7B, the apparatus was composed of an arena with a W30 x L30 x H30 cm “Safe zone” connected to a W15 x L30 x H30 cm corridor (“Interaction zone”) adjacent to a mesh-wire container, in which a rat will be placed in during the test. Test animal was placed into the safe zone with the entrance of interaction zone blocked by an opaque board. A black, opaque shelter (Shaped W10 x L12 x H15 cm, and an opening of W6 x H13 cm on a short edge) were placed at the corner distal to the interaction zone, with the opening facing at the direction in which the entrance of interaction zone was not visible. After 10 minutes of free exploration for acclimatization, a 1 min baseline will be recorded. The rat was then placed into the container, and the opaque board was removed. The animal was allowed to explore both the safe zone and interaction zone for 15 mins.

### Stereotaxic surgery

Surgery was performed at the age of 8-12 weeks for adult heterozygous SF1-Cre mice. Mice were anaesthetized with intraperitoneal (IP) injection of Zoletil/Xylazine mixture, following subcutaneous (SC) injection of ampicillin and carprofen, which prompt analgesics and reduce the chance of infection, respectively. Once the deep anaesthetized state (absence of toe-pinch reflex) was reached, the mouse was transferred to a stereotaxic apparatus and the anesthesia was maintained under 0.3-1.0% isoflurane throughout the procedure. The virus was injected into VMHdm using a pulled glass capillary and an injection pump (RWD) at a flow rate of 1 nL/sec. The same configuration was adopted for virus delivery targeting BNST. The stereotaxic injection coordinates were determined based on the mouse brain atlas^28^ (bregma as origin; VMHdm: medial-lateral ±0.4 mm, anterior-posterior −1.5 mm, dorsal-ventral −5.6 mm; BNST: medial-lateral ±1.0 mm, anterior-posterior 0.05 mm, dorsal-ventral −4.2 mm,)

The optical cannula (0.4 diameter x 7.0 mm length, N.A 0.5, Inper) used for fiber photometry or the GRIN lens (0.6 mm diameter x 7.3 mm length with baseplate attached, Inscopix) used for microendoscopic imaging was lowered to 0.1 mm above the viral injection coordinate and fixed on the skull with Super-Bond C&B and Flow-It™ ALC™. In addition, a protective cover was magnetically affixed to the baseplate. Ampicillin, carprofen, and dexamethasone were administrated subcutaneously daily after surgery for at least 3 days. After surgery, mice were allowed to recover for at least 6 weeks before starting behavioral experiments.

### Major olfactory epithelium (MOE) ablation

To ablate the MOE, test mice were injected intraperitoneally with 3 doses of 2,6-Dichlorobenzonitrile (Dichlobenil, CAS 1194-65-6), an olfactotoxic herbicide, on 7, 5, and 3 days before experiments at a working concentration of 50 mg/mL in 10% DMSO, and a dose of 100 ug/g body weight. Control animals are injected with 10% DMSO alone. Olfactory receptors within the nasal mucosa were expected to be totally ablated by such treatment while leaving vomeronasal organ (VNO) functionally intact^7,10,26^.

### Gonad-removal surgery

Adult BALB/c male/female mice were anesthetized with intraperitoneal (IP) injection of Zoletil/Xylazine mixture, following subcutaneous (SC) injection of ampicillin and carprofen. The anesthesia was maintained under 0.3-1.0% isoflurane throughout the procedure. For male castration, a tiny incision was made on the midline of animal’s scrotum with sterile scalpel. Both testes were subsequently removed. The wound was sutured by surgical skin staple. For female ovariectomy, mouse was first placed in prone position on a heating pad with the lower back shaved and cleaned. A small bilateral dorsolateral incision was made just caudal to the last ribs. Once the muscle layer was dissected and the peritoneal cavity was exposed, the ovary was excised after ligating ovarian pedicle with suture. The wound was then closed with surgical skin staple and sutures. Once the surgery was accomplished, animal was transferred to a clean cage for recovery. Ampicillin and carprofen were administrated subcutaneously daily after surgery for at least 3 days. 2 weeks after surgery, gonad-removed animals were transferred to a clean cage to remove residual sex-hormone-dependent odorants 3 days before being used as social stimulus or intruder.

### Fiber photometry data acquisition

Two LEDs (465 nm and 405 nm) were adopted to excite GCaMP6s through the implanted optical cannula. The 465 nm LED intensity was sinusoidally modulated at 211 Hz and passed through an eGFP excitation filter. The 405 nm LED was modulated at 531 Hz and passed through a 405 nm bandpass filter. Intensities of both LED light sources were separately adjusted in every recording trial to achieve optimal signal-to-noise ratio. Both light streams were coupled to a high NA (0.48), large core (400 mm) optical fiber patch cord, which was attached to a matching implanted optical cannula in each mouse. The 465 nm light excited the GCaMP6s in a Ca^2+^-dependent manner, while 405 nm light that excited GCaMP6s in a Ca^2+^-independent manner, which served as the non-specific reference signals to correct signal artifacts due to photobleaching or animal’s movement. GCaMP6s-emitted fluorescence was collected by the same fiber, passed through a GFP emission filter, and focused onto a photoreceiver and amplifier. Frequency-based lock-in amplifier independently demodulated the emitted fluorescence brightness derived from 405 nm and 465 nm excitation. Behavioral video frame acquisition was synchronized by 20 Hz transistor-transistor logic (TTL) pulses emitted from a digital output port in the photometry console.

### Microendoscopic imaging data acquisition

A head-mounted miniaturized microscope (nVista3, Inscopix) was used for *in vivo* Ca^2+^ imaging. A pilot imaging trial was performed 5 days before behavioral experiments to determine optimal recording parameters, ensuring ideal signal-to-noise ratio, and to minimize photobleaching. Single-cell Ca^2+^ imaging data was acquired at 20 Hz with 2x spatial down sampling, 0.2-0.7 light-emitting diode power, and 1-8x gain, with the parameters determined in accordance with the GCaMP6s expression level indicated by the image histogram displayed in Inscopix Data Acquisition Software (IDAS). Behavioral video frame acquisition was synchronized by transistor-transistor logic (TTL) pulses emitted from the data acquisition box (DAQ).

### Microendoscopic imaging with optogenetic circuit manipulation

To simultaneously perturbate the BNST-VMH pathway with optogenetic approach and capture cellular-resolution Ca^2+^ activities from VMH^SF1^ neurons, mice with opsins expressed in BNST were attached to a head-mounted microendoscope equipped with 620 nm OG-LED (nVoke2.0, Inscopix). To determine how BNST input acutely modulates VMH^SF1^ neurons, mice with ChrimsonR expressed in BNST were placed in an empty home cage, allowing 10 minutes of acclimatization. After 3 minutes of baseline recording, OG-LED stimulation at 5 Hz or 20 Hz was delivered to VMH repetitively in a 5s-on, 10s-off pattern. Light pulse width was set as 5 *μs*, and the power was set at 20 mW/mm^2^. Recording was terminated after 10 minutes of recording. On the other hand, to silence BNST input to VMH during short-term social encounter or social interaction, OG-LED constant light was delivered repetitively to VMH in a 5s- on, 10s-off pattern with 5 mW/mm^2^ power in the presence of introduced conspecific (Silence / BNST-trial).

### Chemogenetic inhibition

Artificial VMH^SF1^ neuron silencing was achieved through “Designer Receptors Exclusively Activated by Designer Drugs” (DREADD)^56^. Six weeks after stereotaxic surgery, mice injected with functional inhibitory DREADD receptor (hM4D(Gi)) or fluorescence control (mCherry) were randomly assigned to CNO-first or Vehicle-first group. Mice in the CNO-first group performed RSI CNO trial on day 1, and RSI Vehicle trial on day 3, and vice versa. Mouse was injected intraperitoneally with CNO (3 mg/Kg) or 10% DMSO in 0.9% saline (Vehicle) and placed into the home cage for the upcoming RSI test. As shown in Figure 8B, the RSI test was initiated 60 minutes post-CNO injection.

### Histology

Mice were deeply anesthetized with Zoletil/Rompun mixture, then subjected to trans-cardiac perfusion with PBS, following by 4% paraformaldehyde (PFA) solution in PBS. Brains were collected and immersed in 4% PFA for more than 1 day, ensuring adequate tissue fixation. Next, the brains were cut into a series of 100-μm coronal sections on a vibratome (5100mz, Campden Instruments). Sections were mounted on Micro slides (PlatinumPro, Matsunami) and sealed using mounting medium with DAPI. The images were captured using epifluorescent microscope (Axioscope A1, Zeiss) or confocal microscope (LSM700, Zeiss) to verify viral expression and optic cannula or GRIN lens placement at 5x or 10x magnification. If the implant tip was positioned outside the target region, or viral expression was absent in the target region, the animal and the corresponding data were excluded from the analysis.

### Quantification and statistical analysis

Data were presented as mean ± standard error of mean (SEM). Sample distribution normality and data homogeneity of variance were first confirmed with D’Agostino– Pearson normality test and Brown-Forsythe test or Bartlett’s test, respectively. Unless otherwise mentioned in figure legends, multi-group comparisons were performed with a one-way analysis of variance (ANOVA) with repeated measures, followed by Bonferroni’s multiple comparison test, the two-tailed paired Student’s t-test was used for paired samples, and the two-tailed unpaired Student’s t-test was used for unpaired samples. Statistical significance was set at *:P < 0.05, **:P < 0.01, ***:P < 0.001 and ****:P < 0.0001. Statistical analyses were carried out using GraphPad Prism 9 and self-written Python scripts.

### Behavioral annotations

Behaviors were manually scored using Behavioral Observation Research Interactive Software (BORIS^29^) in a frame-by-frame manner by trained individuals blinded to synchronized Ca^2+^ signals, and the identities of interacting target. Behaviors were annotated as series of state events; each behavioral event had the onset and termination timing recorded.

For RSI test, we annotated a total of 15 social/non-social behaviors that could be divided into 4 major behavioral categories: Appetitive (approach, sniff/investigation, follow), Defensive (freeze, escape, upright), Consummatory (attack/attack attempt, tail rattle, chase, mount/mount attempt), and Others (allo-groom, self-groom, rear).

For PDA, rat-directed sniff, retrieve (from interaction zone back into shelter), and peek (head or part of body exiting shelter briefly, followed by complete retrieval back into shelter) were annotated as predator-oriented/evoked behaviors. Furthermore, the frame-by-frame positions of test animal were tracked with ezTrack^52^. Whether the test animal was in or out of the shelter was determined by the absence or presence tracking point, with experimenter’s *post-hoc* validation and correction.

### Photometry data preprocessing

The bulk Ca^2+^ dynamics was displayed as ΔF/F ratio, which represented the normalized change in fluorescence signal. ΔF/F Ca^2+^ dynamics were calibrated following the pipeline suggested by Simpson *et al*.^61^ with several personalized modifications. Briefly, fluorescence signal demodulated from GCaMP6s (465 nm excited) or isosbestic GCaMP6s (405 nm excited) were separately resampled to 60 Hz and smoothed using a convolution-based Savitzky-Golay filter with a window size equivalent to 2.5 seconds, and a polynomial order of 3. This filter was operated by successively fitting subsets of adjacent data points with a low-degree polynomial via the linear least square method. Photobleaching of two signals were estimated via fitting a double exponential curve and subtracted from original time series. To estimate the movement-generated artifacts, bleaching-corrected GCaMP6s signals were plotted against bleaching-corrected isosbestic control signals. As movement is assumed to be the major contributor to GCaMP6s signal variations predictable with isosbestic signal, slope and intercept fitted through linear regression were used to generate estimated motion component and subtracted from bleaching-corrected GCaMP6s signal. Least-squares linear fit was subsequently applied to the motion-corrected GCaMP6s signal, producing a fitted F_0_ signal that was used to normalize the GCaMP6s with the equation 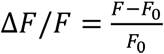.

### Spike detection from photometry bulk Ca^2+^ traces

For bulk Ca^2+^ traces acquired from fiber photometry, spike events with canonical waveform of GCaMP6s Ca^2+^ transient could be identified by performing template search with electrophysiology data acquisition and analysis software (Axon™pCLAMP™ Clampfit 10.7). Searching template was established by averaging 5-10 manually recognized Ca^2+^ transients. Searching threshold was set at 0.1. Identified events went through manual screening process that exclude false events with either noise-like appearance or weak intensity that was indistinguishable from baseline. Accepted events were defined as Ca^2+^ spikes.

### Microendoscopic data preprocessing and ROI extraction

Using Inscopix Data Processing Software (IDPS), Ca^2+^ imaging frames went through 2x spatial down-sampling, spatial band-pass, and motion correction. After the preprocessing, Ca^2+^ activity traces of single neurons were extracted using CNMF-E^18^. Extracted traces and ROIs were carefully inspected to remove duplicate or non-neuronal ROIs. ROIs classified as non-neuronal should exhibit lack of soma-like appearance or artifact-like pattern of corresponding Ca^2+^ traces. To remove Ca^2+^-independent residual noises, Ca^2+^ traces exported from IDPS were smoothed using a convolution-based Savitzky-Golay filter with a window size equivalent to 2.5 seconds, and a polynomial order of 3. Afterwards, the denoised Ca^2+^ trace of each imaged neuron was standardized with the 3-minute (Stimulus exposure test, RSI) or 1-minute (PDA) baseline at the beginning of corresponded recording trial with the formula 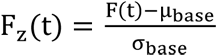, where μ_base_ and σ_base_ reflect the mean fluorescence level and standard deviation over the entire baseline, respectively.

For estimating population averaged Ca^2+^ dynamics in response to external stimulus presentation, the entire Ca^2+^ trace for each imaged neuron acquired from the stimulus exposure test was normalized with formula 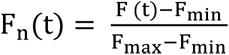, where F_min_^, F^_max_ denote the minimum and maximum values of Ca^2+^ signal from the whole recording trial, respectively. The normalization scaled the entire Ca^2+^ trace to the range of 0 to 1. This effectively normalized the signal-to-noise ratio across neurons from the same animal, preventing strong activation or silencing of a few neurons from dominating the population average features.

### Analyzing neural responses to external stimulus presentation

Multiple stimuli were delivered sequentially in a single trial of stimulus exposure test. To extract stimulus-evoked Ca^2+^ responses from individual neurons or the entire population (fiber photometry), Ca^2+^ signals 10 seconds before to 30 seconds after stimulus introduction were extracted, resampled for length matching, and realigned to the time point of stimulus onset. Extracted stimulus-associated Ca^2+^ dynamics were then z-scored with the 10-seconds pre-stimulus baseline.

The intensity of stimulus-evoked neural response was either presented as post-stimulus peak amplitude or mean activity deviation from pre-stimulus baseline. A neuron would be classified as stimulus-excited if the post-stimulus mean activity increased above 3*σ*, whereas neurons with more than 3*σ* reduction of mean activity would be classified as stimulus-inhibited. Other neurons with activity deviation within the range of ±3*σ* were non-responsive/no-response.

### Analyzing neural response to behaviors

Behavioral data was reduced to point events of behavioral onset, event-associated Ca^2+^ dynamic from individual neuron or the entire population (fiber photometry) was segmented and realigned with a time range of 5 seconds before to 5 seconds after event onset. All the extracted event-associated Ca^2+^ signals were resampled to match the length of template time series, and z-scored with the 5-seconds pre-event baseline. Event-associated z-scored signals were in turn grouped by the type of behavior and directed target and averaged to obtain the mean z-scored Ca^2+^ dynamic of single neuron/population associated to specific behavior. If a behavior occurred less than 3 times in a recording trial, neural response associated with the behavior would be excluded.

The intensity of behavior-associated neural response was either presented as post-stimulus peak amplitude or as mean activity deviation from pre-behavioral baseline. A neuron would be classified as behavior-excited if the maximum post-behavioral activity deviation exceeded +2*σ*, whereas neurons with maximum negative deviation over −2*σ* would be classified as behavior-inhibited. Other neurons with maximum activity deviation within the range of ±2*σ* were non-responsive/no-response.

### Analyzing neural response to BNST-VMH pathway stimulation

Ca^2+^ dynamic from individual neuron was segmented and realigned with a time range of 5 seconds before to 10 seconds after onset of each light pulse delivery. All the extracted event-associated Ca^2+^ signals were resampled to match the length of template time series, and z-scored with the 5-seconds pre-event baseline, and averaged to obtain the mean z-scored Ca^2+^ dynamic of single neuron triggered by light stimulation. A neuron would be classified as light-excited if the mean post-light (only 0 to +5 s included) activity deviation exceeded +2 *σ*, whereas neurons with mean negative deviation over −2*σ* would be classified as light-inhibited. Other neurons with maximum activity deviation within the range of ±2*σ* were non-responsive/no-response.

### Preference score

The strength of neuronal preference for either one of the targets from a selected pair of stimuli was defined as 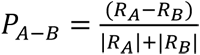, where R_A_ and R_B_ represented the response intensity for stimulus A and B, respectively. Neurons with P_A-B_ = 1 might either be exclusively excited by stimulus A, exclusively inhibited by B, or excited by A and inhibited by B, and vice versa.

### Choice probability

Choice probability (CP) is used to determine whether a single neuron might be probabilistically tuned to either of two events. To calibrate the CP of a neuron for a pair of conditions, histograms of the neuron’s Δ*F*/*F*_0_under each of the two conditions were plotted against each other, generating a receiver-operating characteristic (ROC) curve. The CP value is the integral of area under such ROC curve. CP value shall lie within the range of 0 to 1, with CP of 0.5 implying absence of difference between two conditions.

To determine if a neuron’s CP is statistically significant. We calibrated the CP of the same neurons with the bout timings of the two conditions shuffled. Shuffling was repeated for 100 times, from which we estimated the mean and s.t.d of shuffled CPs. A neuron is considered tuned to a specific behavior or stimulus if it exhibits a CP of less than 0.3 or greater than 0.7 and ≥ 2*σ* deviation from the shuffled mean.

### Identifying stimulus-specific neurons

To determine the breadth of tuning for each neuron, we utilized its stimulus-triggered responses to calibrate normalized entropy with the formula^34,62^:

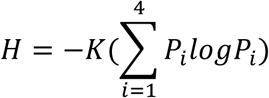

Where *P*_*i*_ represents the proportional response of a neuron to each of the four stimuli and *K* is a scaling constant that constrain *H* in the range of 0 to 1. Neurons with broad tuning to multiple stimuli tend to have higher *H* values. On the other hand, *H* = 0 signifies that the neuron exclusively responded or not responded to just one stimulus. We also calibrate the noise-to-signal ratio for each neuron to compare the magnitude of response across stimuli. The noise-to-signal ratio was calibrated via dividing the proportional response to the second-best stimulus (noise) by the proportional response to the best stimulus (signal)^36^. Higher noise-to-signal ratio implies that the neuron responded in similar level to at least two stimuli, while lower noise-to-signal ratio signifies a large contrast between maximum stimulus-triggered response to the others.

We defined neurons with entropy less than 0.5 as “Stimulus selective”, which was further divided into two subgroups by noise-to-signal ratio cutoff at 0.5. Stimulus-selective neurons with noise-to-signal ratio less than 0.5 were characterized as showing positive specificity. These neurons were selectively activated by one stimulus with the other three stimuli triggering substantially weaker responses. In contrast, stimulus-selective neurons with noise-to-signal ratio above 0.5 were said to had negative specificity. These neurons were activated at a comparable level by three stimuli, while specifically silenced by or not responsive to the remained stimulus

### Generalized linear model

To predict neural activity from behavior and external stimulus, we trained generalized linear models to describe the Ca^2+^ dynamic of each neuron as a weighted linear combination of multiple behaviors and stimulus presence with the formula:

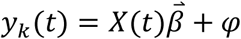

Where *y*_*k*_(*t*) reflects the Ca^2+^ activity of neuron *k* at time *t*, 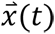 denotes a feature matrix composed of binary vectors representing the presence of specific external stimulus or occurrence of certain behavior, 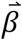 is a behavior-filter describing how a neuron integrates information over time, and the error term is *φ*. The model was fitted using 10-fold cross validation with non-negative least-angle regression (LARS) employment of least absolute shrinkage and selection operator (LASSO)^6^. The performance of model fitting is reported as cross-validated *R*^2^ (*cvR*^2^). Neurons with all available regressors explaining less than 10% variance were categorized as residuals.

### Hierarchical clustering

Ca^2+^ traces were subjected to principal component analysis (PCA) along time dimension. The number of kept PCs was determined via the cumulative explained variance (until total explained variance exceed 80%). The values of PCs in all neurons are subjected to agglomerative hierarchical clustering utilizing Euclidean distance and Ward’s-linkage. The optimal number of clusters is determined by elbow method for KMeans clustering.

### Decoder analysis

To test if stimulus identities are decodable from population neural activities, we established linear Support Vector Machine (SVM) decoders trained with imaging frames in which each interested stimulus was presenting. The trained decoders are in turn tested with imaging frames during stimulus presentation recorded from a held-out trial. For each mouse, decoding was repeated 20 times, with the decoder’s performance in discriminating stimulus identities based on imaged activities reported as the averaged F1 score across repetition. To evaluate the significance of stimulus decoding, performance of “Data” decoder trained on observed stimulus labels was compared to “Shuffled” decoder trained on shuffled data. The shuffle was repeated 100 times and averaged to yield the F1 score of Shuffle decoder.

For behavioral decoding, similar training and performance evaluation procedure were utilized. Nevertheless, data labels reflecting behavioral presence/absence or discrepant behavioral conditions are not likely to be balanced due to sparseness of some behaviors or irregular expression of behavior across imaging session. Therefore, equal number of frames corresponded to distinct behavioral conditions were included for decoder training, ensuring that the chance level decoder performance will be 0.5.

### Dimensionality reduction

Low-dimensional representations that aid visualizing time-evolving population neural dynamics were constructed using PCA. Whereas disparities in stimulus-triggered Ca^2+^ activity of each neuron for distinct external cues are visualized by performing UMAP embedding.

### Cosine similarity

To estimate the level of discrepancy in neural trajectory associated with different conditions, we adopt Cosine similarity as a metric for similarity measurement, which is calibrated via the formula:

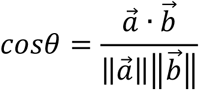

Where 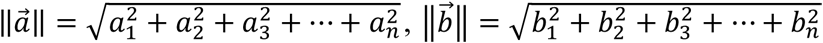, and 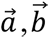 denote the mean vector of neural trajectory corresponded to condition a and b, respectively. A cosine similarity of 0 indicates that the vectors are orthogonal (unrelated), while a cosine similarity of 1 implies that two vectors are perfectly aligned. In contrast, a cosine similarity value of −1 reflects that the vectors are perfectly dissimilar (opposite). Recorded population activity is first subjected to PCA along neural dimension. Limited number of the first few PCs that collectively explain more than 80% of total variance are included to yield the mean neural vectors.

