## Supplementary figures and legends for "A Ventromedial Hypothalamic Neuron Subset Encodes a Conspecific-Tuned Behavior State Driving Social Investigation"

### Supplementary Figure Legends

**Figure S1. Stimulus representation of VMH$^{SF1}$ neurons is stable across days, related to Figure 1 and 2.**

**(A)** Male-triggered $Ca^{2+}$ response from a representative subgroup of neurons with similar dynamics across days.

**(B)** Heatmaps showing the male-induced $Ca^{2+}$ response of stimulus-responsive neurons across 3 recording sessions.

**(C)** Histogram showing the distribution of Pearson's correlation coefficient for stimulus-evoked $Ca^{2+}$ traces across recording sessions. Vertical dashed lines mark the population-averaged PCC.

**(D-E)** Percentage of registered neurons showing same type of stimulus-triggered response across 2 recording sessions (D) or all 3 sessions (E).

**(F)** Composition of neurons with consistent stimulus-triggered response across recording sessions.

**(G)** Composition of neurons with inconsistent stimulus-triggered response across recording sessions. M: Male mouse; F: Female mouse; Obj: Inanimate object; R: Rat.

**Figure S2. Decodable and selective stimulus representation of VMH$^{SF1}$ neurons, related to Figure 2.**

**(A-B)** Acute stimulus-evoked $Ca^{2+}$ dynamics of stimulus-specific neurons in **Figure 2J**.

**(C)** Confusion matrix showing the stimulus decoding performance of SVM decoder trained on real (left) or shuffled (right) data.

**(D)** Accuracy of data-trained SVM decoder in decoding stimulus identity. (n = 16 mice; One-sample Student's t-test, hypothesized population mean = 25).

**Figure S3. Fiber photometric assessment of the sensory modality vital for maintaining male-biased conspecific sex preference of VMH$^{SF1}$ neurons, related to Figure 3.**

**(A)** Example photos of an anesthetized (top) or a dangled awake (bottom) male mouse.

**(B)** Bulk $Ca^{2+}$ responses of VMH$^{SF1}$ neurons to an awake, anesthetized, or dangled male mouse.

**(C-D)** Peak $\Delta F/F$ of male- triggered populational activity.

**(E)** Schematic illustration of experimental treatment for selective removal of olfactory pathway. MOE: Major olfactory epithelium; VNO: Vomeronasal organ.

**(F)** Bulk $Ca^{2+}$ dynamics triggered by male/female mouse with/without functional MOE.

**(G)** Discrimination factor of male/female mouse induced bulk$Ca^{2+}$ response before/after MOE ablation, defined as the peak amplitude of male- triggered divided by female-triggered responses.

**(H)** Averaged $\Delta F/F$ deviation of male- triggered populational activity before/after MOE ablation.

**(I)** Bulk $Ca^{2+}$ dynamics triggered by a GI male, CM, or female mouse.

**(J)** Discrimination factor of male/female-induced bulk $Ca^{2+}$ response with/without pheromonal inputs from male mouse.

**(K)** Averaged $\Delta F/F$ deviation of male- triggered populational activity with/without pheromonal inputs.

**Figure S4. VMH$^{SF1}$ neurons respond with reversed sex preference to investigation targeting gonad-removed conspecifics, related to Figure 3 and 5.**

**(A)** Same as **Figure 5A**, but with gonad-removed conspecifics.

**(B)** Choice probability histogram and percentages of neurons tuned to male-or female-directed investigation with gonad-intact (left) or gonad-removed (right) conspecifics introduced.

**(C)** Correlation of population mean response to male versus female investigation with gonad-intact (left) or gonad-removed (right) conspecifics introduced.

**(D)** ROIs from a representative mouse showing the relative abundance and the spatial distribution of neurons with distinct responsiveness for investigation events directed toward different conspecific sexes.

**(E)** Population-averaged variance explained by full encoding model or simple regression models. (n = 11 mice).

**(F)** Fraction of neurons significantly modulated by each regressor.

**(G)** Behavior-associated population neural trajectories in common subspace spanned by the first 3 PCs, corresponded to CM/OVXF-directed investigation.

**(H)** Explained variance of the first 5 PCs of sniff male/female response.

**(I)** Performance of SVM decoder in predicting CM/OVXF investigation.

**Figure S5. Silencing BNST-VMH pathway renders VMH$^{SF1}$ neurons more tuned to female presence without altering investigation tuning, related to Figure 4 and 5.**

**(A)** Illustration of light delivery pattern during social interaction in sham/silence trial.

**(B)** Heatmaps showing the averaged z-scored response of individual neurons in sham (left) or silence (right) RSI trial to investigation events targeting conspecifics of different sexes. $Ca^{2+}$ dynamics of neurons were sorted by the magnitude and direction of post-behavioral mean $Ca^{2+}$ level deviation.

**(C)** Choice probability histogram and percentages of neurons tuned to male- or female-directed investigation with (left) or without (right) BNST-VMH pathway activity.

**(D)** Population-averaged variance explained by full encoding model or simple regression model of specified stimulus/behavioral events in sham or silence trial. (n = 4 mice).

**(E)** Fraction of neurons significantly modulated by each regressor in sham (left) or silence (right) trial.

**(F)** Performance of SVM decoder in predicting male/female investigation from held-out neural data in sham (left) or silence (right) trial.

**Figure S6. VMH$^{SF1}$ neuronal response to social investigation versus consummation or defense, related to Figure 5 and 6.**

**(A)** Experimental procedure and setup for fiber photometric measurement of $VMH^{SF1}$ population $Ca^{2+}$ dynamics during reciprocal social interaction (RSI) test.

**(B)** A representative bulk $Ca^{2+}$ trace during social interaction.

**(C)** Fraction of large bulk $Ca^{2+}$ spikes (> 3 $\sigma$ above baseline) temporally associated with specified social behavioral categories.

**(D)** Behavioral-averaged bulk $Ca^{2+}$ trace.

**(E)** Behavioral-averaged bulk activity deviation in response to specified behavioral category (One-sample Student's t-test, hypothesized population mean = 0).

(**F-I**) Fraction of all neurons classified as behavior-excited (EXC), -inhibited (INH) or non-responsive (NR) to (F) male-directed investigation/attack (n = 652 neurons from 9 mice), (G) male-directed investigation/escape (n = 474 neurons from 6 mice), (H) male-directed investigation/mount (n = 619 neurons from 7 mice), or female-directed investigation/mount (n = 599 neurons from 7 mice).

**(J)** (left) Population-averaged $Ca^{2+}$ trace of neurons aligned with the onset of male-directed investigation/mount. (right) Population-averaged response amplitude to male-directed investigation/mount. (n = 7 mice)

**(K)** Population-averaged investigate versus mount male preference scores (One-sample Student's t-test, hypothesized population mean = 0.5).

**(L)** Choice probability histogram and percentages of neurons tuned to male-directed investigation or mount. (n = 619 neurons from 7 mice).

**(M-O)** Same as **J-L**, but for female-directed investigation versus mount. (n = 599 neurons from 7 mice)

**Figure S7. Behavior-associated Ca$^{2+}$ dynamics of VMH$^{SF1}$ neurons during predator defense, related to Figure 7.**

**(A)** A sample behavioral and neural data showing how shelter transition events were identified and used for calibrating transition-associated Ca$^{2+}$ activity of single neuron (right).

**(B)** Choice probability histogram and percentages of neurons preferring in-shelter or out-shelter (n = 997 neurons from 15 mice)

**(C)** Choice probability histogram and percentages of neurons preferring shelter-exit or -entry (n = 997 neurons from 15 mice)

**(D)** Population mean Ca$^{2+}$ deviation associated to shelter-exit/entry.

**(E)** Behavioral shelter preference plotted against sniff-rat-triggered population activity deviation.

**(F)** Heat maps of individual neuron averaged Ca$^{2+}$ dynamics (top) aligned with onset of predator-triggered behaviors. Neurons were sorted independently by the timing of peak Ca$^{2+}$ activity for each behavior. Population-averaged Ca$^{2+}$ dynamics are showed in the bottom panels.

**Figure S8. Combinatorial VMH$^{SF1}$ neural responses to social investigation and shelter transitions under predatory threats, related to Figure 7.**

**(A)** The Ca$^{2+}$ activities from RSI and PDA tests are normalized with mean and standard deviation of trial-associated baseline period, merged, and subject to agglomerative hierarchical clustering. Each row is color-coded according to the neuron's corresponding cluster and the preference class it belongs to.

**(B)** ROIs from a representative mouse showing the relative abundance and the spatial distribution of neurons with distinct responsiveness for shelter transition in PDA or female-directed social investigation.

**(C)** Percentage of neurons that are selectively responsive or co-responsive to shelter transition or female-directed social investigation. (n = 7 mice)

**(D-E)** Fraction of neurons showing distinct responsiveness to shelter exit and male-(D) or female-directed (E) investigation (left), with detailed composition of combinatorial response pattern for dual-responsive neurons (Both) showed in the right panel.

**(F-G)** Same as **D-E**, but for shelter entry versus mouse investigation.

### Figure S1.

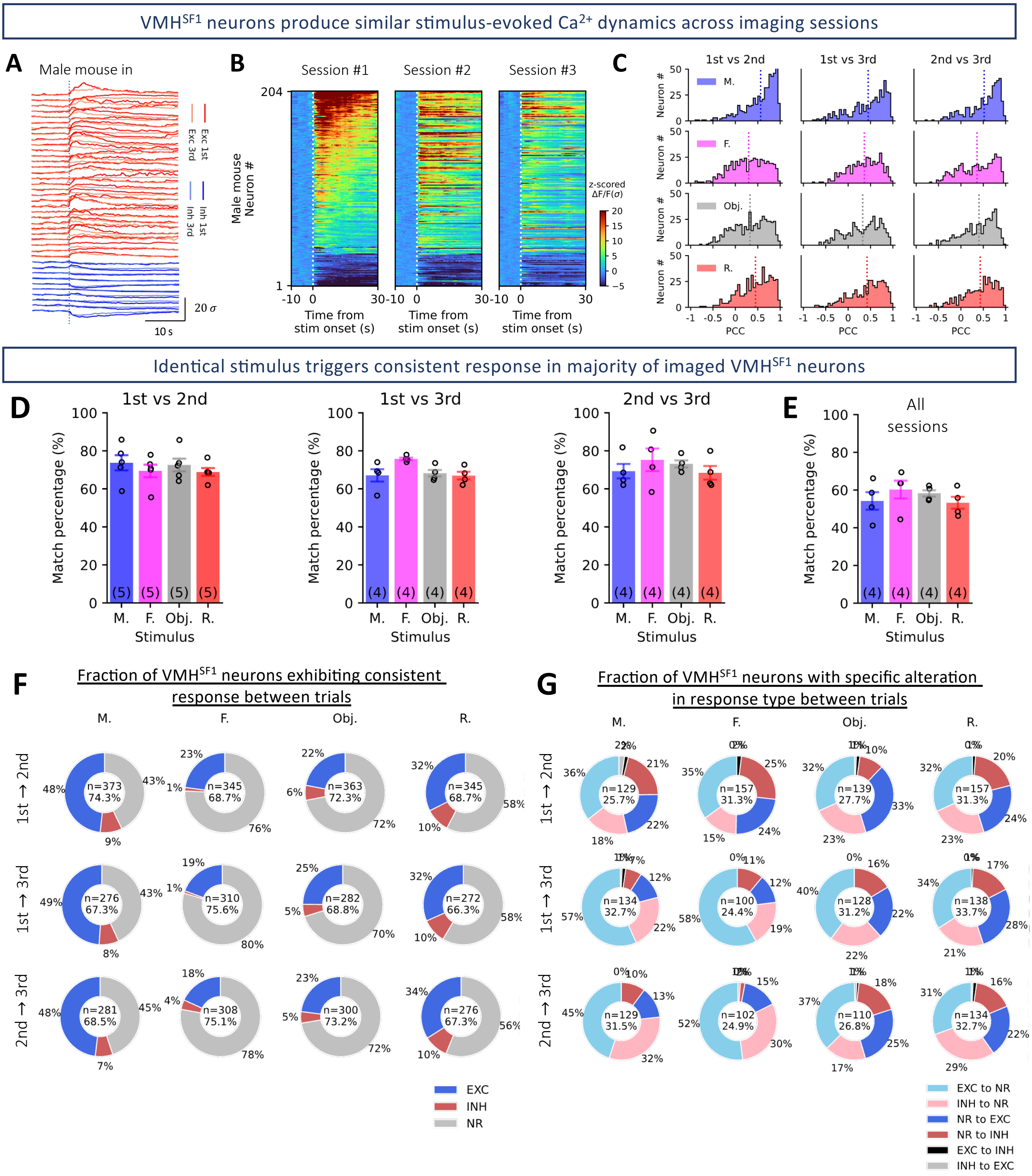

VMHSF1 neurons produce similar stimulus-evoked Ca2+ dynamics across imaging sessions
A Male mouse in
B Session #1 Session #2 Session #3
Male mouse Neuron #
Time from stim onset (s)
z-scored ΔF/F(σ)
C 1st vs 2nd 1st vs 3rd 2nd vs 3rd
Neuron #
PCC
Identical stimulus triggers consistent response in majority of imaged VMHSF1 neurons
D 1st vs 2nd 1st vs 3rd 2nd vs 3rd
E All sessions
Match percentage (%)
Stimulus
F Fraction of VMHSF1 neurons exhibiting consistent response between trials
G Fraction of VMHSF1 neurons with specific alteration in response type between trials
EXC
INH
NR
EXC to NR
INH to NR
NR to EXC
NR to INH
EXC to INH
INH to EXC

### Figure S2.

Detailed stimulus-evoked $Ca^{2+}$ dynamics of stimulus-specific $VMH^{SF1}$ neurons

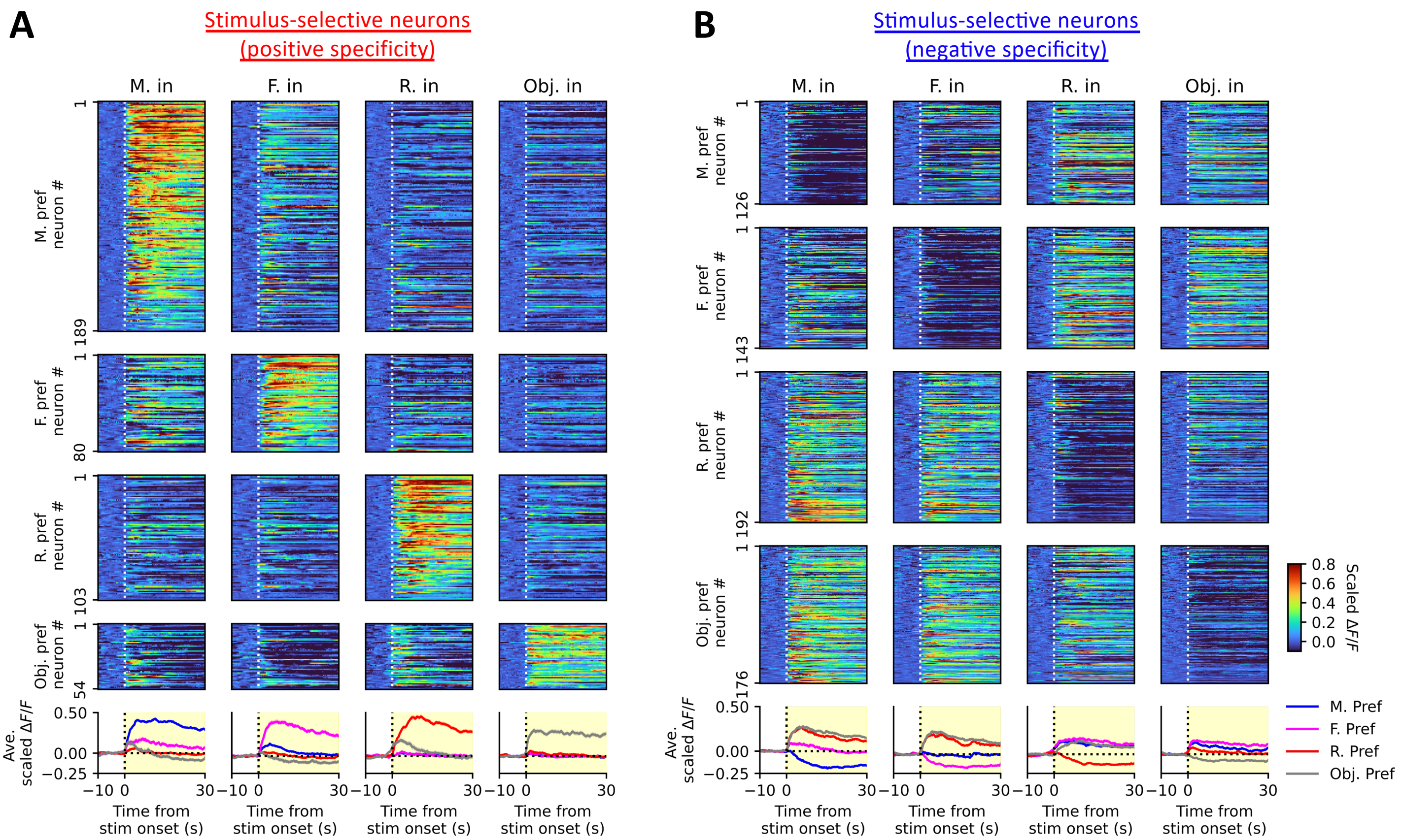


$VMH^{SF1}$ neurons form long-term, decodable stimulus representation for social and non-social cues

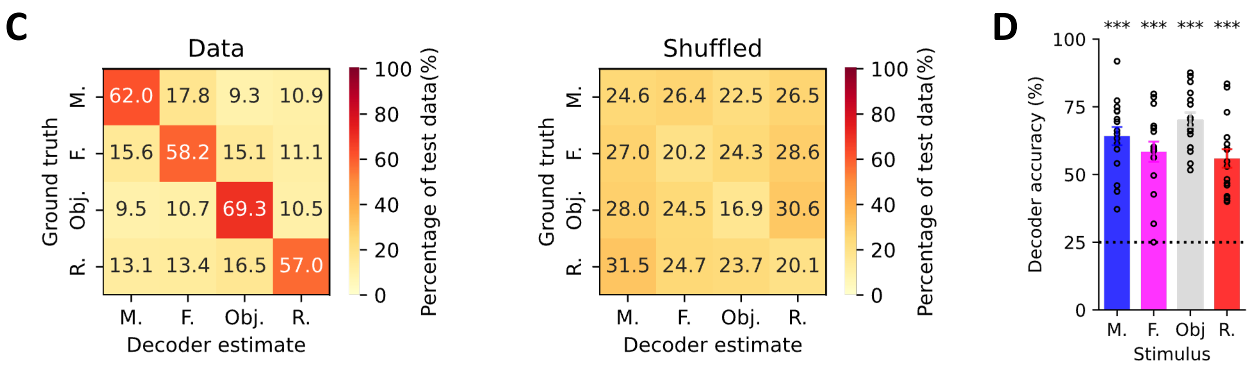

### Figure S3.

Inaccessible social cue evokes weaker VMH$^{SF1}$ populational response

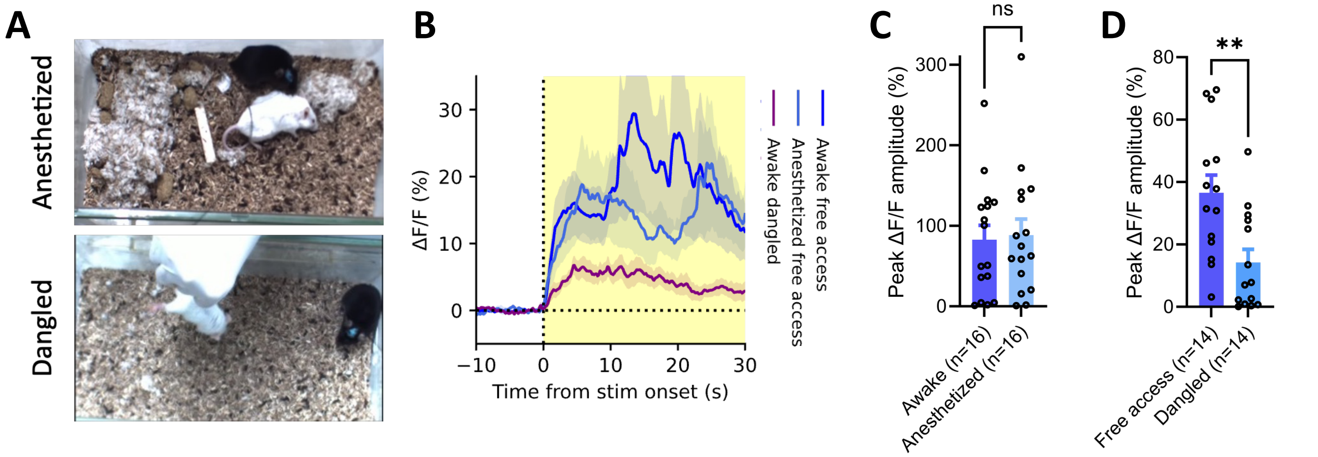


VMH$^{SF1}$ populational male preference requires VNO-, but not MOE-relayed olfactory signals

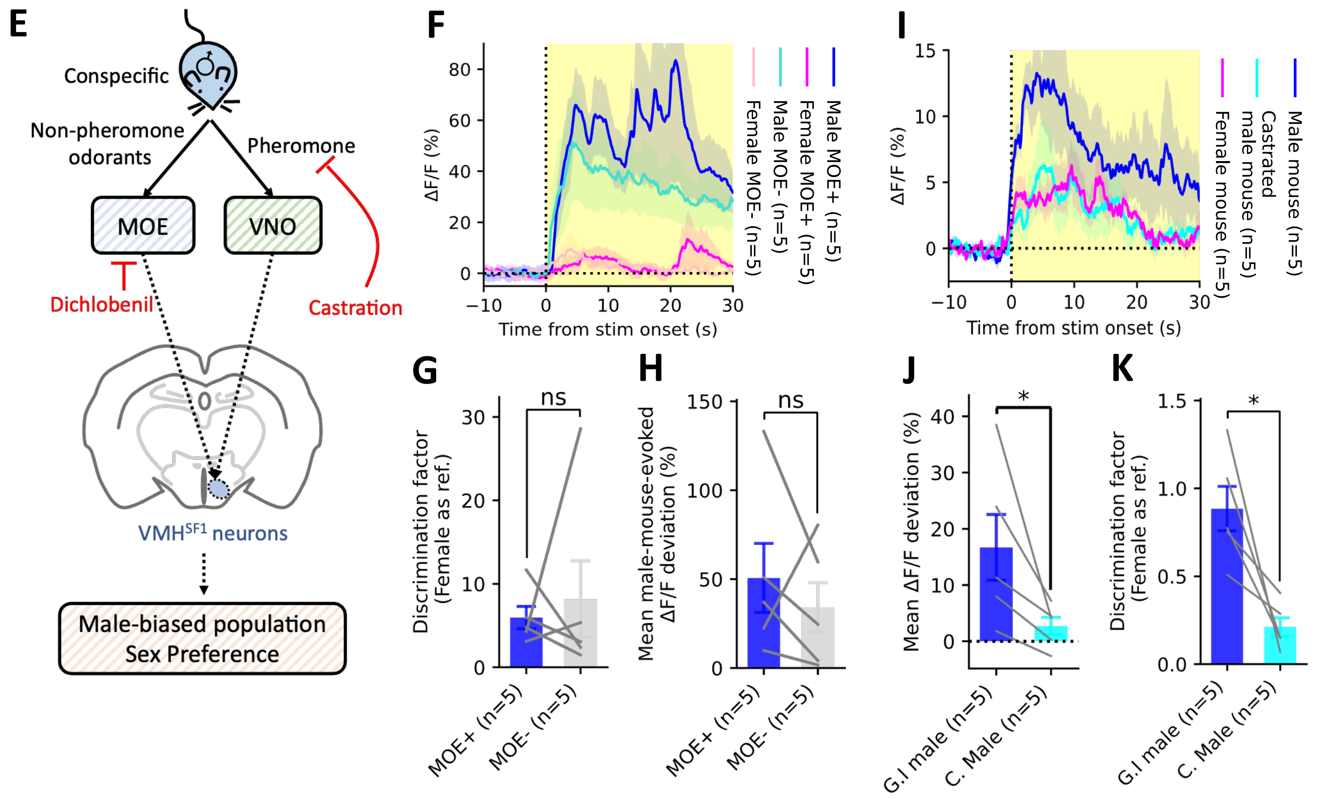

### Figure S4.

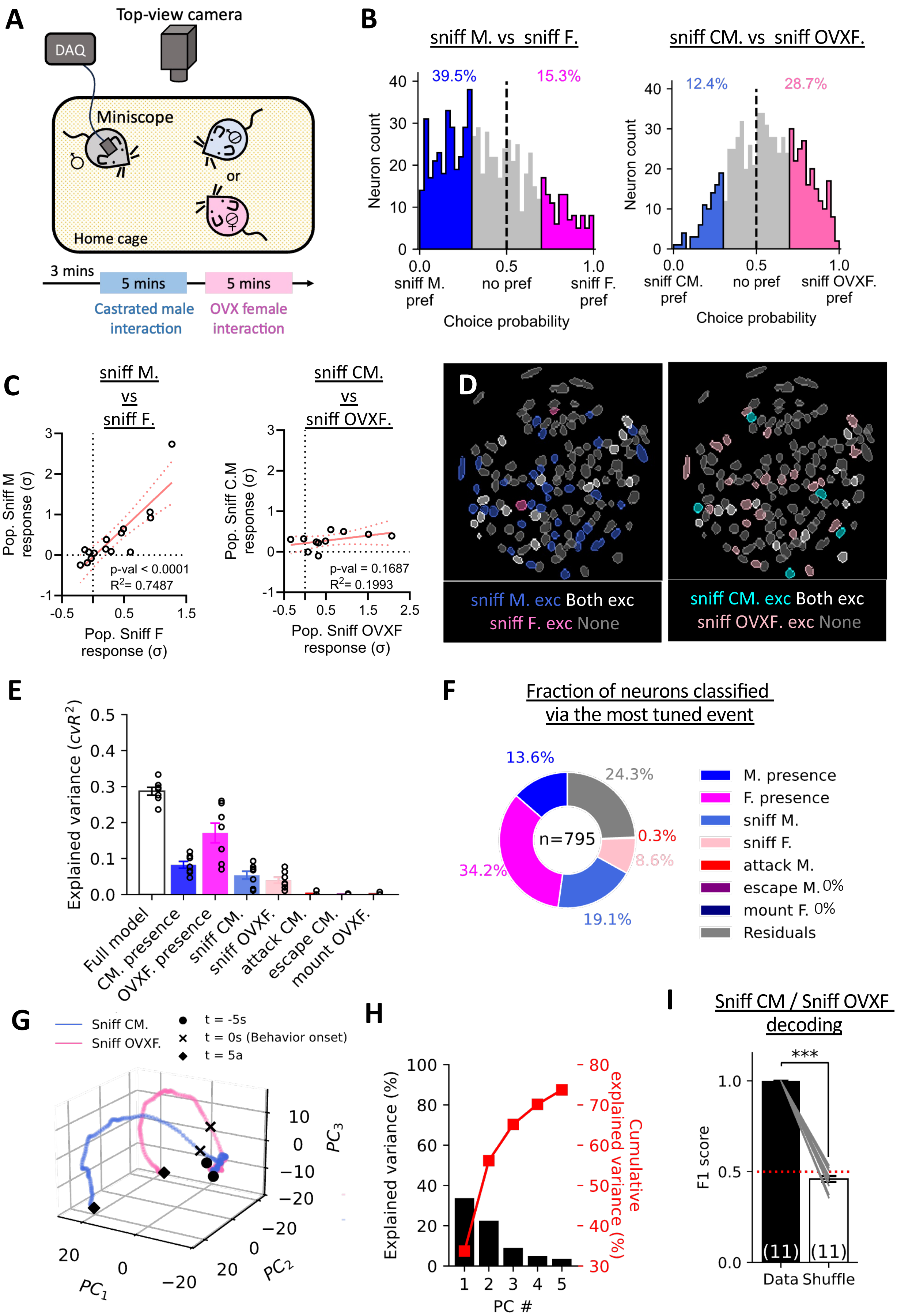

A
Top-view camera
DAQ
Miniscope
or
Home cage
3 mins
5 mins
5 mins
Castrated male interaction
OVX female interaction
B
sniff M. vs sniff F.
39.5%
15.3%
Neuron count
0.0 sniff M. pref
0.5 no pref
1.0 sniff F. pref
Choice probability
sniff CM. vs sniff OVXF.
12.4%
28.7%
0.0 sniff CM. pref
0.5 no pref
1.0 sniff OVXF. pref
C
sniff M. vs sniff F.
Pop. Sniff M response (σ)
p-val < 0.0001
R2= 0.7487
Pop. Sniff F response (σ)
sniff CM. vs sniff OVXF.
Pop. Sniff C.M response (σ)
p-val = 0.1687
R2= 0.1993
Pop. Sniff OVXF response (σ)
D
sniff M. exc Both exc
sniff F. exc None
sniff CM. exc Both exc
sniff OVXF. exc None
E
Explained variance (cvR2)
Full model
CM. presence
OVXF. presence
sniff CM.
sniff OVXF.
attack CM.
escape CM.
mount OVXF.
F
Fraction of neurons classified via the most tuned event
13.6%
24.3%
0.3%
8.6%
34.2%
19.1%
n=795
M. presence
F. presence
sniff M.
sniff F.
attack M.
escape M. 0%
mount F. 0%
Residuals
G
Sniff CM.
Sniff OVXF.
t = -5s
t = 0s (Behavior onset)
t = 5a
PC1
PC2
PC3
H
Explained variance (%)
Cumulative explained variance (%)
PC #
I
Sniff CM / Sniff OVXF decoding
***
F1 score
(11)
(11)
Data
Shuffle

Figure S5.

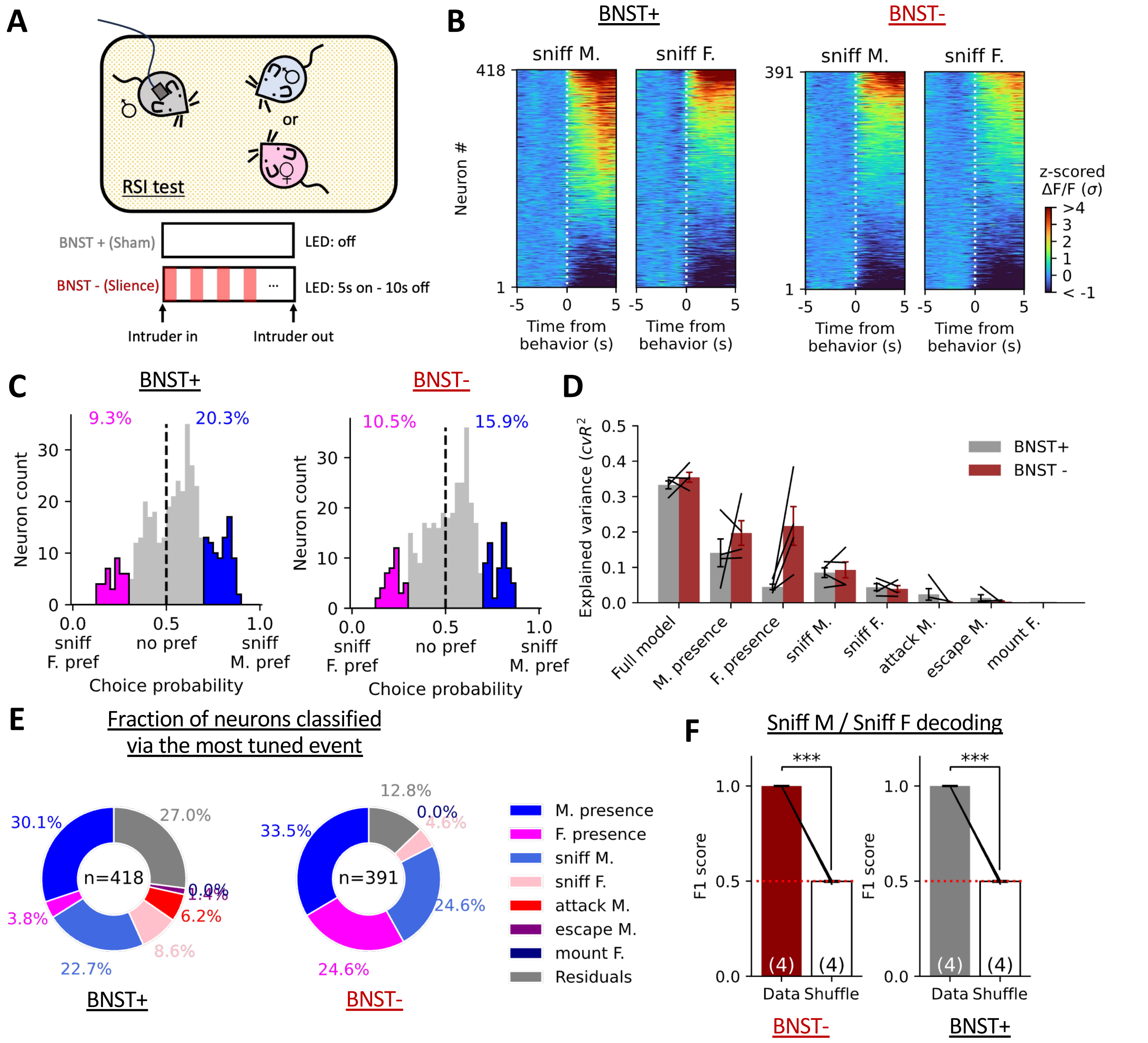
A
RSI test
or
BNST + (Sham)
LED: off
BNST - (Slience)
LED: 5s on - 10s off
Intruder in
Intruder out
B
BNST+
BNST-
sniff M.
sniff F.
Neuron #
Time from behavior (s)
z-scored ΔF/F (σ)
C
BNST+
BNST-
9.3%
20.3%
10.5%
15.9%
Neuron count
sniff F. pref
no pref
sniff M. pref
Choice probability
D
Explained variance ($cvR^2$)
BNST+
BNST -
Full model
M. presence
F. presence
sniff M.
sniff F.
attack M.
escape M.
mount F.
E
Fraction of neurons classified via the most tuned event
n=418
n=391
30.1%
27.0%
0.0%
1.4%
6.2%
8.6%
22.7%
3.8%
12.8%
0.0%
4.6%
33.5%
24.6%
24.6%
M. presence
F. presence
sniff M.
sniff F.
attack M.
escape M.
mount F.
Residuals
BNST+
BNST-
F
Sniff M / Sniff F decoding
F1 score
***
(4)
Data
Shuffle
BNST-
BNST+

### Figure S6.

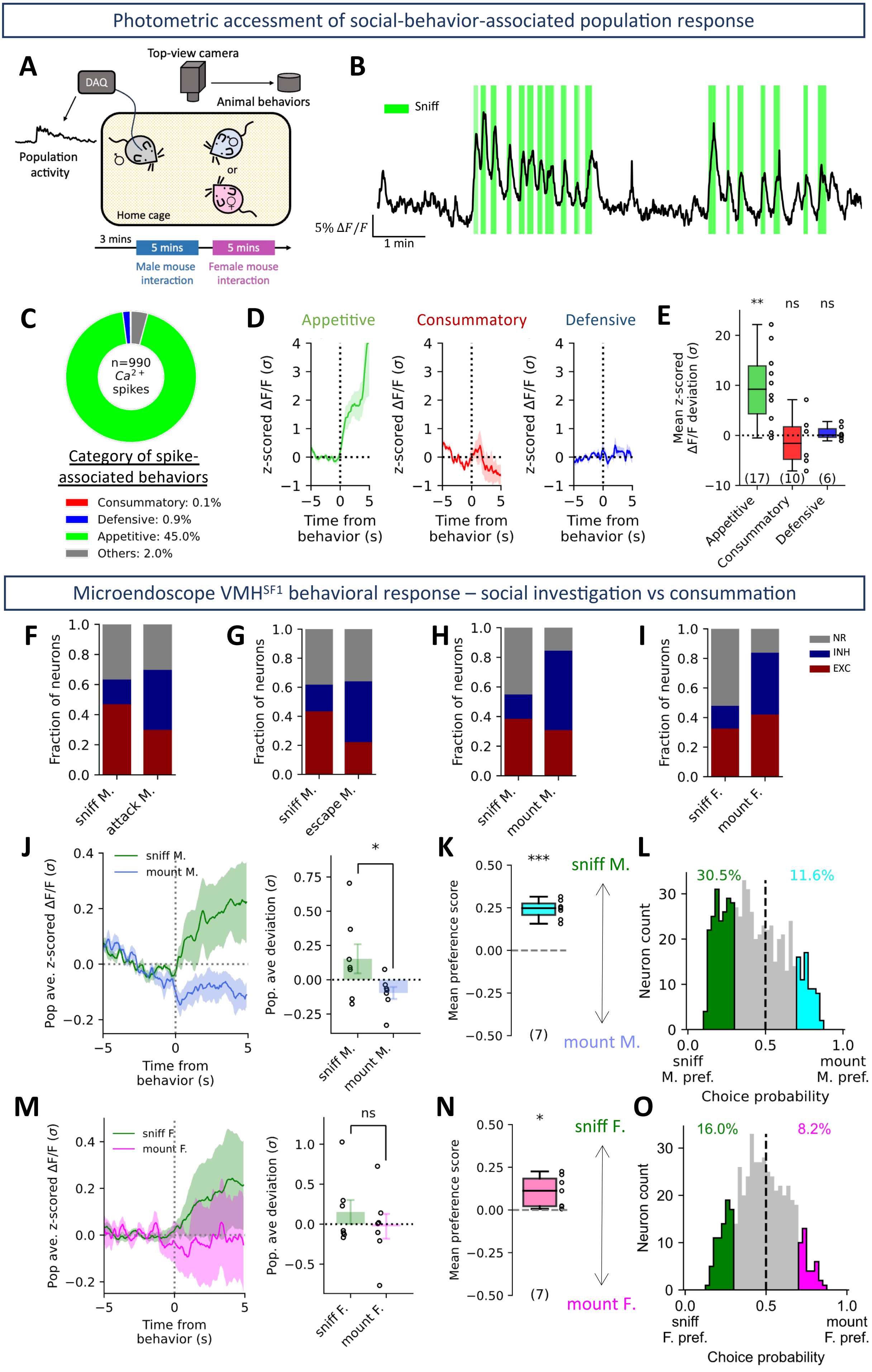
Photometric accessment of social-behavior-associated population response
A
Top-view camera
DAQ
Animal behaviors
Population activity
or
Home cage
3 mins
5 mins
5 mins
Male mouse interaction
Female mouse interaction
B
Sniff
5% ΔF/F
1 min
C
n=990
Ca2+
spikes
Category of spike-associated behaviors
Consummatory: 0.1%
Defensive: 0.9%
Appetitive: 45.0%
Others: 2.0%
D
Appetitive
Consummatory
Defensive
z-scored ΔF/F (σ)
Time from behavior (s)
E
Mean z-scored ΔF/F deviation (σ)
**
ns
ns
(17)
(10)
(6)
Appetitive
Consummatory
Defensive
Microendoscope VMHSF1 behavioral response – social investigation vs consummation
F
G
H
I
Fraction of neurons
NR
INH
EXC
sniff M.
attack M.
sniff M.
escape M.
sniff M.
mount M.
sniff F.
mount F.
J
sniff M.
mount M.
Pop ave. z-scored ΔF/F (σ)
Time from behavior (s)
Pop. ave deviation (σ)
*
K
***
sniff M.
mount M.
Mean preference score
(7)
L
30.5%
11.6%
Neuron count
sniff M. pref.
mount M. pref.
Choice probability
M
sniff F.
mount F.
Pop ave. z-scored ΔF/F (σ)
Time from behavior (s)
Pop. ave deviation (σ)
ns
N
*
sniff F.
mount F.
Mean preference score
(7)
O
16.0%
8.2%
Neuron count
sniff F. pref.
mount F. pref.
Choice probability

### Figure S7.

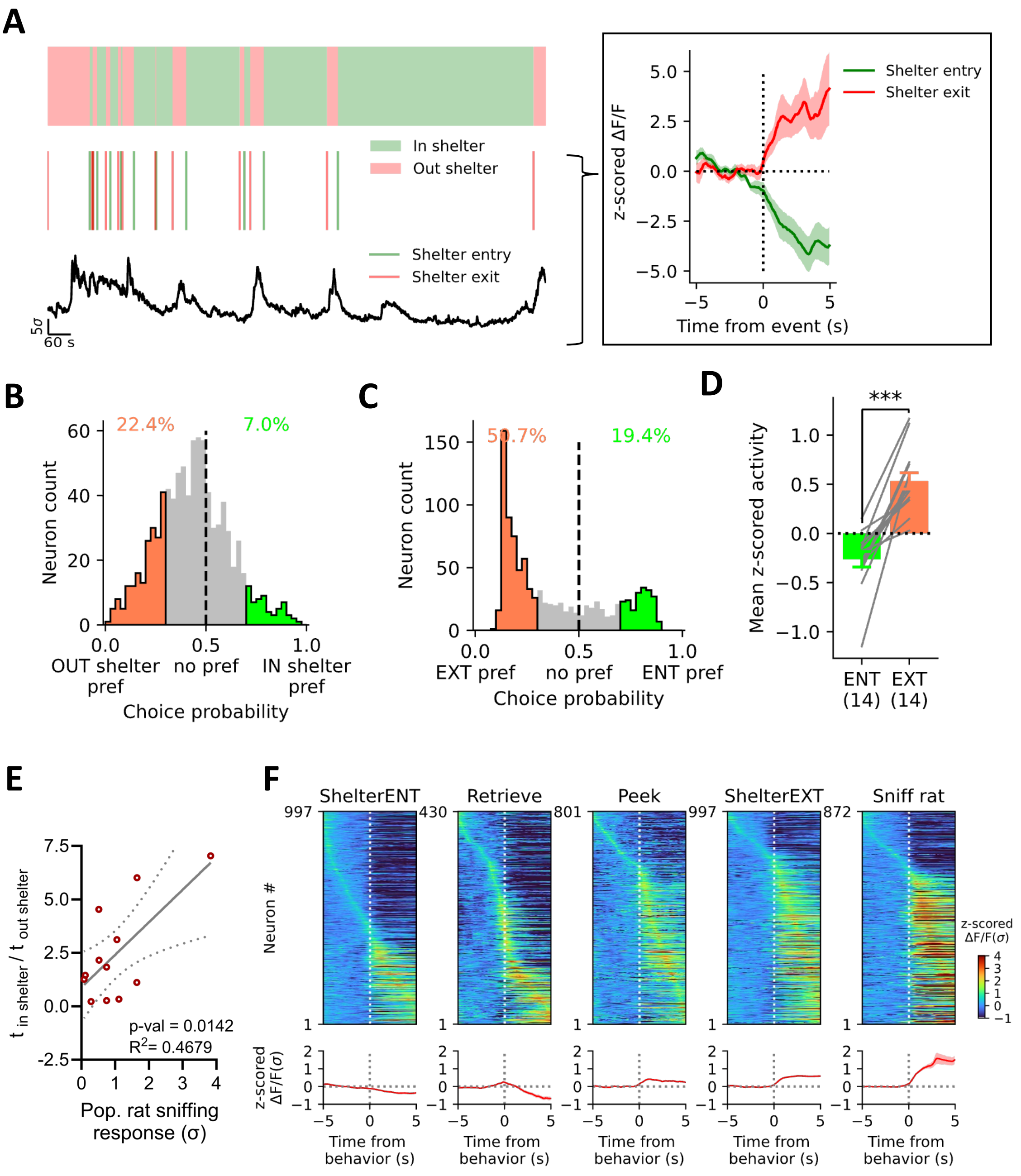

A
In shelter
Out shelter
Shelter entry
Shelter exit
5σ
60 s
z-scored ΔF/F
Time from event (s)
B
22.4%
7.0%
Neuron count
OUT shelter pref
no pref
IN shelter pref
Choice probability
C
50.7%
19.4%
Neuron count
EXT pref
no pref
ENT pref
Choice probability
D
***
Mean z-scored activity
ENT (14)
EXT (14)
E
t in shelter / t out shelter
p-val = 0.0142
R2= 0.4679
Pop. rat sniffing response (σ)
F
ShelterENT
Retrieve
Peek
ShelterEXT
Sniff rat
Neuron #
z-scored ΔF/F(σ)
Time from behavior (s)

Figure S8.

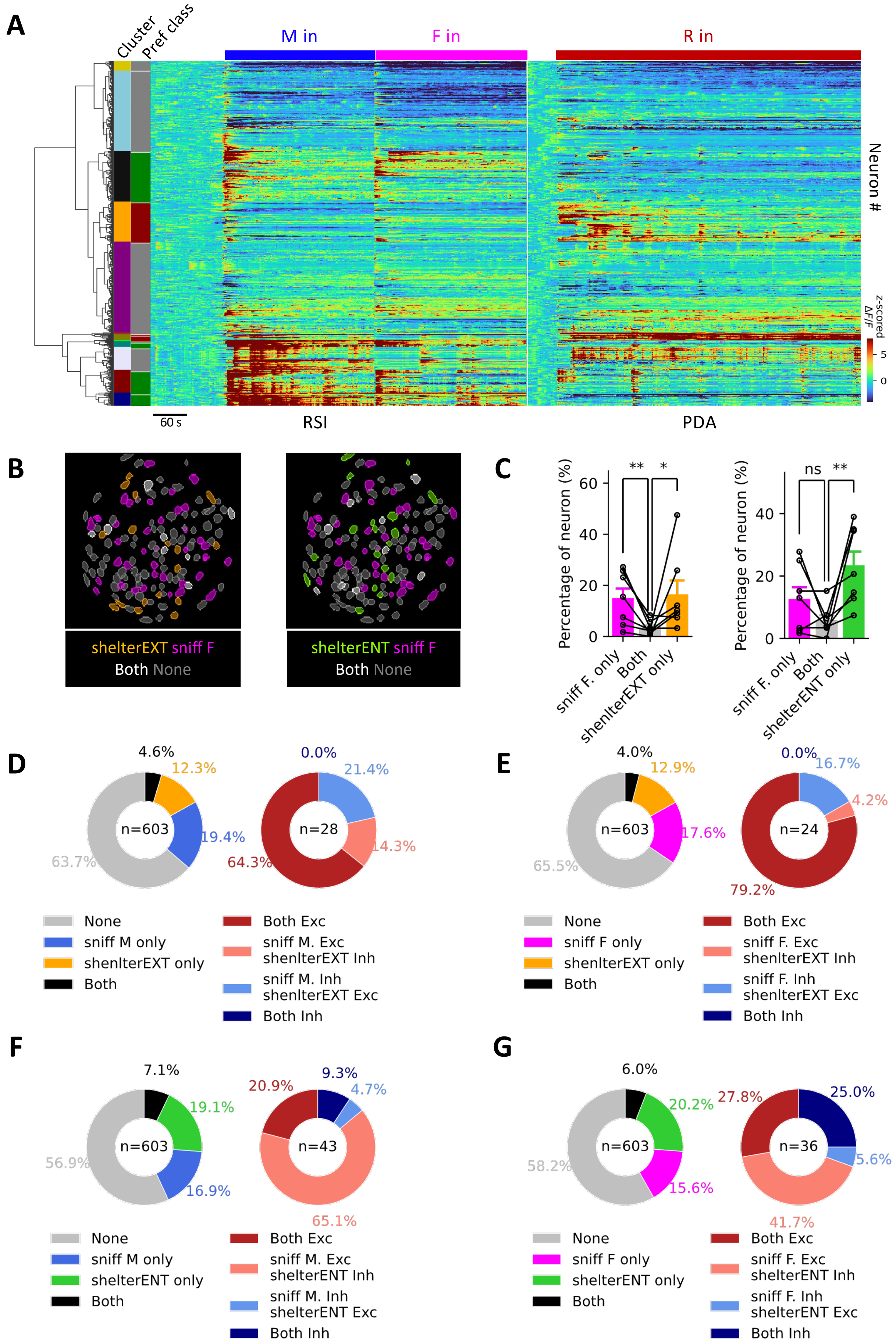

A
Cluster
Pref class
M in
F in
R in
Neuron #
z-scored ΔF/F
5
0
60 s
RSI
PDA
B
shelterEXT sniff F
Both None
shelterENT sniff F
Both None
C
Percentage of neuron (%)
60
40
20
0
**
*
sniff F. only
Both
shenlterEXT only
ns
**
sniff F. only
Both
shelterENT only
D
4.6%
12.3%
19.4%
63.7%
n=603
0.0%
21.4%
14.3%
64.3%
n=28
None
sniff M only
shenlterEXT only
Both
Both Exc
sniff M. Exc shenlterEXT Inh
sniff M. Inh shenlterEXT Exc
Both Inh
E
4.0%
12.9%
17.6%
65.5%
n=603
0.0%
16.7%
4.2%
79.2%
n=24
None
sniff F only
shenlterEXT only
Both
Both Exc
sniff F. Exc shenlterEXT Inh
sniff F. Inh shenlterEXT Exc
Both Inh
F
7.1%
19.1%
16.9%
56.9%
n=603
20.9%
9.3%
4.7%
65.1%
n=43
None
sniff M only
shelterENT only
Both
Both Exc
sniff M. Exc shelterENT Inh
sniff M. Inh shelterENT Exc
Both Inh
G
6.0%
20.2%
15.6%
58.2%
n=603
27.8%
25.0%
5.6%
41.7%
n=36
None
sniff F only
shelterENT only
Both
Both Exc
sniff F. Exc shelterENT Inh
sniff F. Inh shelterENT Exc
Both Inh
